# Contrasting temperature-induced gene network rewiring and isoform switching underlie relative thermal tolerance of coral species

**DOI:** 10.64898/2025.12.26.696607

**Authors:** Federica Scucchia, Pierrick Harnay, Hollie M Putnam

## Abstract

Organismal thermal tolerance will shape future coral reef diversity and abundance, as ocean warming drives extensive bleaching and mortality through increasingly frequent marine heatwaves. Coral thermal tolerance varies markedly among species, reflecting differences in physiology underpinned by molecular regulatory mechanisms. Gene expression plasticity enables transcriptional adjustments to environmental stress, while alternative splicing and isoform switching further expand molecular adaptability by altering transcript diversity and abundance. Because genes function within combinatorial networks, changes in these network members and in the identity of highly influential hub genes can reveal higher-order biological reorganization under stress. Yet, the contribution of isoform switching to gene network dynamics remains unknown in corals. Here we integrate physiology, gene network topology, hub genes dynamics, and isoform switching across three Hawaiian corals, *Porites compressa*, *Montipora capitata*, and *Pocillopora acuta,* to identify molecular processes shaping thermal performance. We show that the interplay between expression plasticity, hub gene rearrangement, and isoform switching drives species-specific thermal tolerance. Network analysis uncovered extensive rewiring and temperature-responsive hub genes turnover regulated by switches in isoform usage. The most thermally tolerant species, *P. compressa*, shows high energy reserves, stable constitutive gene expression and continuous isoform adjustments across temperatures, while fine-tuning thermal response through dedicated hub genes. In contrast, the most thermally sensitive species, *P. acuta,* shows lowest energy reserves, with broad transcriptomic shifts and isoform switching controlling the reorganization of network leadership at high temperatures. Together, these findings identify isoform switching as a newly recognized regulatory mechanism contributing to coral thermal resilience and species persistence in warming oceans.

## Introduction

Ocean warming driven by climate change has caused widespread coral mortality and is among the most urgent threats to the survival of coral reefs today (1, 2). With the steady rise in mean ocean surface temperatures, marine heatwaves are occurring more often and lasting longer (3, 4). These extreme warming events are a primary driver of coral bleaching, a stress response caused by the collapse of coral-algal symbiosis (5) that is becoming more frequent and widespread (6, 7), contributing to the loss of ∼50% of live coral cover over the last 75 years (8). In the coral-algal nutritional symbiosis, corals acquire the majority of their energy from the algal partners (9, 10); hence, sustained bleaching leads to depletion of carbon supply as well as the coral host’s energy reserves (11–13), which is increasingly driving vast coral mortality (1, 14, 15). Mass loss of live coral rapidly erodes the structural framework of the reef (16, 17), leading to deterioration of ecosystem services and goods that range from coastal protection to fisheries support and tourism (18).

Not all coral species show the same responses to increase water temperature, as bleaching susceptibility varies depending on host and symbiont characteristics including colony size (19–21), microbiome community profiles (22, 23), host tissue biomass (14, 24), genetic variability within symbiont types (25–28) and host genomics (29–32). In recent years, the role for coral host gene expression plasticity and regulation in acclimatization to environmental stress has emerged (33–35), indicating that regulatory mechanisms, both genetic and epigenetic in nature (36), are key to the relative tolerance of corals in response to temperature stress.

Alternative splicing (AS), a mechanism of gene expression regulation by which a pre-mRNA sequence produces different transcript isoforms from the same gene, increases proteome diversity without changing the underlying genomic sequence (37). AS is an important post-transcriptional regulatory event during response to temperature stress in plants (38–41), crustaceans (42), tunicates (43), oysters (44), fish (45–48) and in corals (49). The recent study characterizing AS patterns in corals in response to heat stress demonstrates that nearly half (40%) of the protein coding genes in *Acropora cervicornis* display AS (49), which indicates AS is a critical yet understudied regulatory mechanism for thermal stress response in stony corals (49). Beyond single species studies, it is therefore critical to determine how the use of differentially spliced isoforms relates to varying thermal thresholds displayed by different coral species. In other sessile taxa, i.e. plants, AS induced by heat stress regulates cross-species acclimatization to different thermal thresholds, mediated by changes in the relative abundance of particular isoforms (reviewed in (50)). Isoform switching, or differential isoform usage, represents a central element of the eukaryotic adaptive toolkit, enabling cells to respond to abrupt or sustained temperature changes, for example by altering protein stability, subcellular localization, or regulatory interactions, and thereby directly shaping cellular protection and stress resilience (40, 51).

Cells can also respond to stress and environmental stimuli through dynamic re-organization of gene regulatory networks, one of the most important organizational levels within the cell where signals are integrated in terms of changes in gene expression (52–54). Within a gene network, genes with the most interactions are defined as hub genes. These genes act as network leaders, coordinating the expression of multiple pathways and thereby exerting disproportionate control over cellular stress responses (53, 55–60) and adaptation to different environments (52, 53). Elucidating hub genes dynamics and isoform switching through AS induced by stress can greatly improve our understanding of cellular states. Indeed, hub genes are important regulators of AS and isoform abundance (61), and integrating isoform switching with hub genes patterns has uncovered how post-transcriptional regulation reshapes cellular pathway coordination and functional output (62). However, the interplay between these two regulatory layers has been largely understudied in stony corals. Thus, integrative approaches that combine thermal performance assessments, physiological traits, hub genes expression and isoform switching analyses are critically needed to unravel the mechanisms underlying coral thermotolerance.

Here, we tested the thermal sensitivity of three dominant reef-building coral species in Hawai’i, *Pocillopora acuta*, *Montipora capitata* and *Porites compressa,* using an integrative assessment of holobiont physiological functions, temperature-induced gene networks rewiring of hub genes and isoform switching patterns. To establish species-specific thermal tolerance, we first generated photosynthesis-irradiance curves across six temperatures, between 12°C to 35°C (12, 18, 25, 26.8, 30 and 35°C). We modeled thermal performance curves to quantify the temperature dependence of key coral performance rates (i.e. photosynthesis and respiration), and assessed the underlying host and symbiont physiological parameters. We further examined gene expression plasticity, expression breakpoints, and gene network functional re-organization (i.e., shifting hub genes). Lastly, we tested the temperature-dependent dynamics of alternative splicing and differential isoform usage relative to the hub genes. The powerful methodological framework we employed allowed us to identify the biological signatures, from cell to whole organism, of coral thermal tolerance, and revealed both network rewiring and isoform switching as previously unexplored mechanisms in corals. These insights into the multi-level regulation of coral thermal tolerance advance fundamental understanding of coral resistance and resilience, providing critical knowledge into the future trajectory of reef ecosystems in a dramatically changing ocean.

## Results and Discussion

### Divergent thermal performances and physiological underpinnings

Coral bleaching is driven by the combination of high light and temperature (63, 64), thus quantifying the photosynthesis versus irradiance curves (PI curves) across a range of temperatures generates essential information for bleaching outcomes. Thermal performance can be quantified by a set of metrics extracted from curve fitting of thermal performance curves (TPCs; (65)). These include the minimum (*ctmin*), maximum (*ctmax*), and optimum (topt) temperature of performance; rates of activation (*e*) and deactivation (*eh*), maximum rate (*rmax*), and rate at a reference temperature (*rtref*), as well as the range of temperatures over which the curve’s rate is at least 80% of peak rates (*breadth*). Species-specific thermal tolerances are clearly revealed by PI parameters (Fig. S1, S2, Tables S1, S2) and TPCs (Fig. 1, Fig. S3 and Table S3). *P. compressa* showed superior performance across most photosynthetic TPC metrics than *M. capitata* and *P. acuta* (*ctmax, breadth, rmax, rtref* and *eh*; Fig. 1B, Fig. S3, Fig. S4 and Table S3). In contrast, *M. capitata* exhibited the greatest performance for *ctmin*, *e* and *topt*, whereas *P. acuta* did not show superior performance for any of the photosynthesis metrics (Fig. 1B, Fig. S3 and Table S3). Respiratory performance followed a similar trend, with *P. compressa* displaying the greatest performance for most TPC metrics (*ctmin* and *el*; Fig. 1D, Fig. S3 and Table S3). Species-specific thermal *breadth* and *topt* values indicate that *P. compressa* behaves as a generalist species, whereas *M. capitata* and *P. acuta* exhibit specialist capacities (65–69). With a higher *topt*, *M. capitata* and *P. acuta* may be favored, at least initially, with increases in seawater temperature. However, further temperature rise will likely be detrimental, as indicated by their higher *eh* and lower photosynthesis to respiration ratio at high temperature as compared to *P. compressa* (Fig. S3 and S5). The latter supports *P. compressa* as the best in photosynthetic performance, particularly at high temperatures (Fig. S5).

**Fig. 1.**
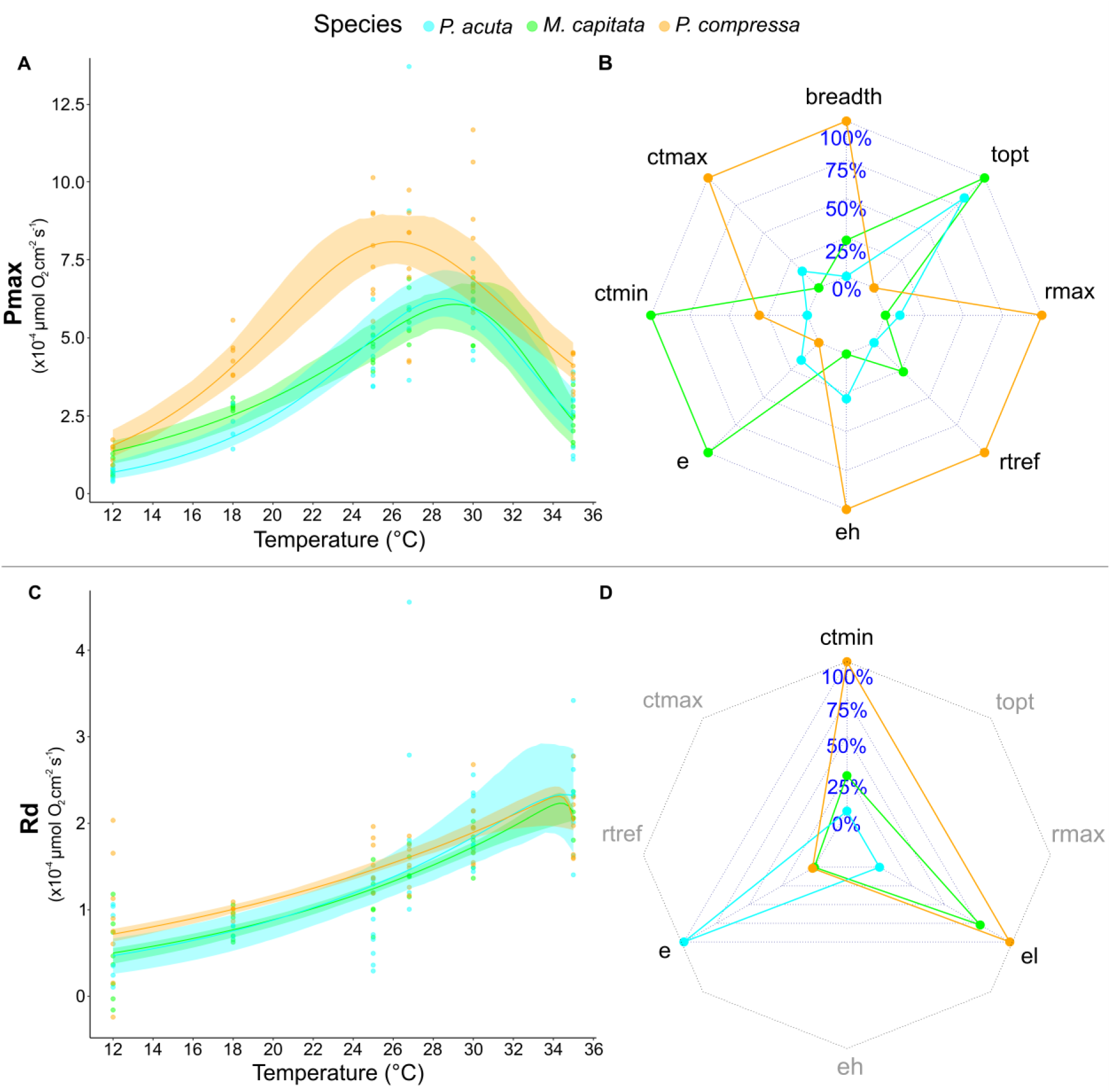
Thermal performance of photosynthesis and respiration rates across species. Fitted TPCs for each PI-derived parameter (maximum photosynthetic rate in A and respiration in C) are shown with 95% confidence intervals. TPC-generated metrics (Fig. S3) were standardized and plotted using a radar plot for (B) Pmax and (D) Rd, where each axis represents a standardized metric and each species’ thermal profile is shown as a colored polygon. In (B) and (D), metrics are directionally oriented so that for metrics where higher values indicate greater thermal tolerance (e.g., *ctmax, topt, breadth, rmax, rtref*), standardized values were used as-is; for metric where lower values indicate greater tolerance (e.g., *e, eh, el, ctmin*), the sign of the standardized value was reversed, to ensure that higher values on all axes consistently represent greater thermal tolerance.

Since TPCs vary across biological organization levels (70), it follows that such generalist-specialist trade-offs involve different underlying physiological mechanisms. Baseline physiological traits at the control temperature revealed that *P. compressa* reaches significantly higher values of biomass and protein concentrations than the other species (Fig. S6 and Tables S4, S5), consistent with a thicker tissue, which provides protection for symbiont communities as well as enhanced energy reserves (14, 71–73). Energetic resources, such as carbohydrates, are critical to maintain ATP levels during periods of thermal stress (74), enhancing coral resistance to bleaching (71, 75–77) and supporting larger coral respiratory demands and energetic requirements (78, 79). Indeed, *P. compressa* has larger carbohydrates concentration than *P. acuta* (Fig. S6, Tables S4, S5), and appears to possess higher lipid concentration compared to *M. capitata* (80), indicating that a larger energetic pool is available for this species, influencing its ability to better support respiratory demands under increasing temperature (Fig. 1D). The superior energetic status of *P. compressa* is also in part driven by its higher symbiont density and chlorophyll *a* and *c* concentrations compared to both *M. capitata* and *P. acuta*, observed at the control temperature and across most other temperature treatments (Fig. S7, S8, Tables S6, S7).

Given the short duration of exposure, we did not expect strong temperature-driven changes in algal physiology, and indeed, cell density and chlorophyll concentrations remained relatively stable across temperatures within each species (Fig. S9-11), indicating that differences among species primarily reflect baseline physiological state. Baseline differences in the physiological traits and states of corals generate differences in how they perform under stress (81), which has been hypothesized to be driven by energy reserves (14) but also by environmental preconditioning (73). We therefore asked whether baseline physiological differences are reflected in distinct molecular response trajectories under thermal stress.

### Thermal stress triggers species-specific transcriptomic trajectories

Transcriptomic analyses revealed that 35°C is the temperature treatment that elicits the strongest expression changes in all coral species, with *P. acuta* also responding at 30°C (Fig. S12). Global variation in gene expression at the highest temperatures (30°C and 35°C) among the three species, explored through discriminant analysis of principal components (DAPC) and Markov chain Monte Carlo (MCMC) mixed models, indicates that magnitude of gene expression plasticity was similar at 30°C across the three species, but it differed markedly at 35°C (Fig. 2). *P. acuta* displayed the greatest plasticity, with a 100% posterior MCMC-based probability of greater plasticity than both *P. compressa* and *M. capitata*. For *P. acuta*, the responses to 30 °C and 35 °C occurred in the same direction along the LD1 axis, implying reliance on similar expression patterns across moderate and high thermal stress. In contrast, *M. capitata* and *P. compressa* exhibited shifts in opposite directions on LD1 at 30°C as compared to 35 °C, which suggests a hormetic response where the temperature beyond the organismal threshold triggers the engagement of different expression patterns. Between species at 35°C, the direction and magnitude of *P. acuta*’s transcriptomic plasticity was opposite to the other two species, pointing to fundamentally different thermal response mechanisms.

**Fig. 2.**
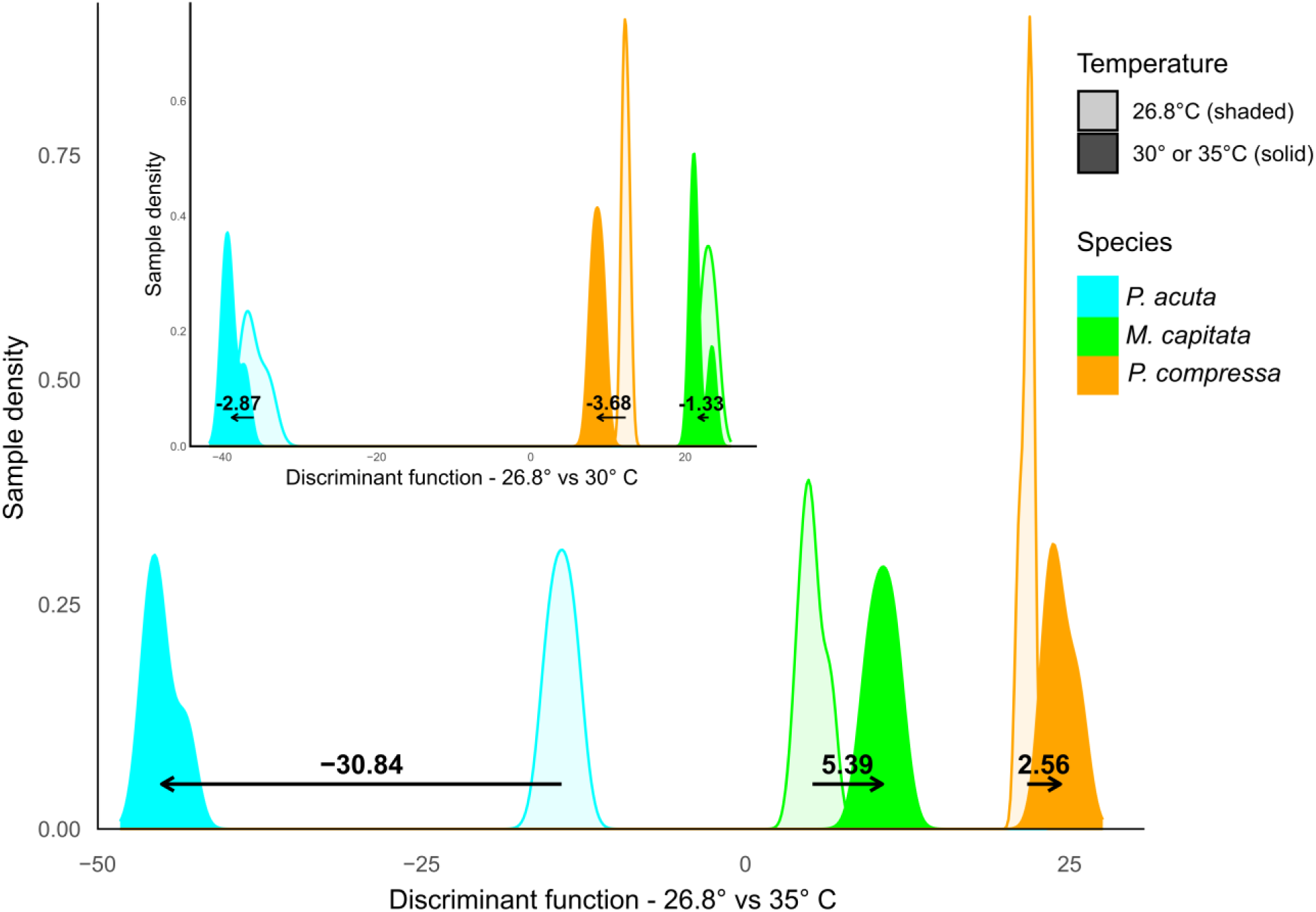
Variation in expression plasticity at high temperature across species. The first linear discriminant (LD1) scores were plotted as density curves for each species, with different temperature treatments distinguished by transparency levels (control samples shown with reduced opacity, and 30/35°C samples shown as solid lines). Mean LD1 scores were calculated for control and elevated temperature treatments within each species and plotted as arrows to indicate the direction and magnitude of the shift in gene expression from control to elevated temperature conditions (arrow length proportional to the plasticity response). Numerical labels above arrows display the difference in mean LD1 scores. Color indicates species (green = *M. capitata*, cyan = *P. acuta*, orange = *P. compressa*).

Further, we found that the gene expression profile of *P. compressa* and *M. capitata* at the control temperature is much different than the one of *P. acuta* (distance between the peaks on x axis in the 26.8°C versus 30°C comparison in Fig. 2), indicating that different expression patterns and/or different magnitude of expression characterize the transcriptomic signatures at ambient conditions. This, combined with *P. acuta* showing a much stronger shift of expression, thus higher plasticity than the other two species at 35°C, indicates a diverse set of high baseline expression (*P. compressa* and *M. capitata*) versus high plasticity (*P. acuta*) mechanisms employed by the different species. Such conditioned or dampened responses in the more tolerant corals suggests differential temperature sensing mechanisms compared to the more sensitive species (e.g., Transient Receptor Potential (TRP) channels or their copy number (82, 83)), or more likely acclimatory gene regulatory mechanisms due to different internal tissue conditions (84). In fact, the higher internal heterogeneity and large physicochemical gradients of thicker tissues (85, 86) may pre-condition thick tissue corals to withstand environmental perturbation through environmentally-mediated priming (73, 87). If this is indeed the case, thick-tissued species and with perforate skeletons such as *P. compressa* and *M. capitata* (88), where the tissue reaches much deeper layers that imperforate corals as *P. acuta* (10), would be primed and better acclimated to external environmental variability. If the internal environment of thin-tissue corals is more homogenous and more closely reflects external conditions, environmental perturbations would elicit a strong response (73, 87), as in the case of *P. acuta.* Whereas in the thick-tissue corals *P. compressa* and *M. capitata* an increased constitutive level of gene expression (i.e., frontloading, (34)) dictated by conditioning exposure to wide internal fluctuations would reduce the transcription requirement for inducible stress responses.

Notably, the gene expression shift of *P. compressa* and *M. capitata* at 35°C is in the opposite direction compared to the shift at 30°C (Fig. 2), resembling a dampening response. Both plasticity and dampening represent alternative routes to enhance coral thermal tolerance (33, 34, 89–91). Transcriptional dampening, which is associated with reduced plasticity, can result from a combination of transcriptomic resilience and/or frontloading, which can provide tolerance to stress through faster return to baseline levels (92). At the same time, a larger capacity for plasticity may replace elevated baseline expression in conferring higher stress tolerance (92, 93). However, transcriptomic plasticity and a high baseline expression of stress response pathways may not confer resilience at the holobiont level, given that the symbiotic algae might not have the same thermal tolerance capacity, as shown by the higher photosynthetic temperature sensitivity of *P. acuta* and *M. capitata* at 35°C (Fig. S5).

### Gene networks reorganization through shifts of hub genes at high temperatures

Network preservation and hub gene dynamics revealed distinct molecular architectures underlying species-specific thermal tolerance, investigated using weighted gene co-expression network analysis. We found that *P. acuta* displays a broader response to temperature stress, with two large modules (Fig. S13) encompassing a wide set of genes showing significant break in pattern at the highest temperatures (30°C and 35°; (Fig. 3A and Table S8)). Whereas *P. compressa* and *M. capitata* show a more streamlined or targeted response, with regulatory changes distributed across multiple smaller gene networks (Fig. S14 and S15), specifically 4 *M. capitata* module clusters (Fig. 3A, Fig. S16 and Table S8) and 5 *P. compressa* module clusters (Fig. 3A, Fig. S16 and Table S8). Overall, for all species, most clusters show a significant break-point between the control and the 35°C treatment (Fig. 3A), in accordance with the strongest gene expression response that we observed at the 30°C and 35°C temperature treatments (Fig. S12).

**Fig. 3.**
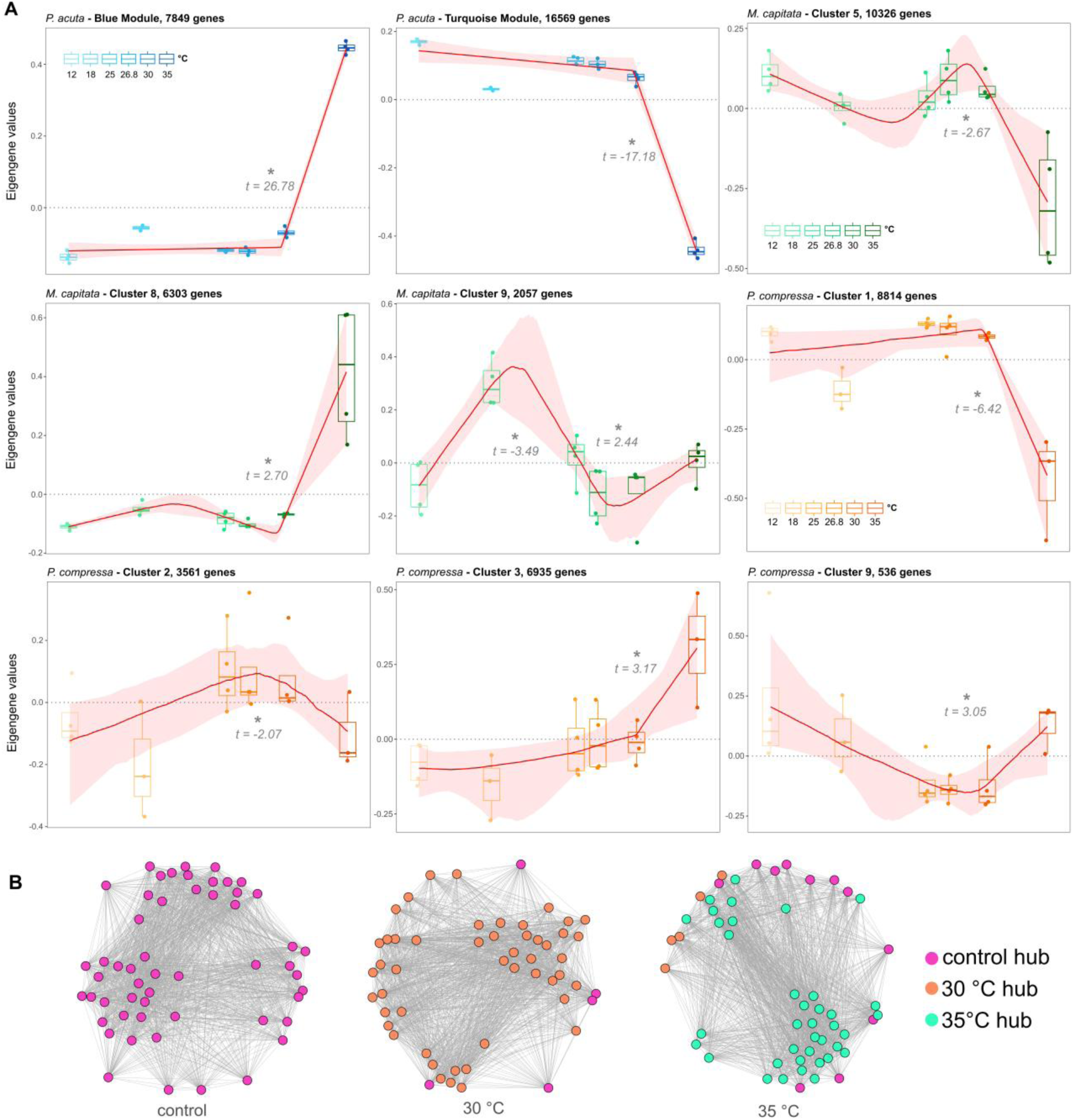
Within-species dynamic changes in gene expression profiles and gene network rewiring across temperatures. A) WGCNA clusters/modules with significant break-points at higher temperatures than control. The variation in eigengene expression of clusters with significant temperature correlations was analyzed to detect break-points (change-points) in the relationship between temperature and the eigengene value. Two modules for *P. acuta*, three clusters for *M. capitata*, and 4 clusters for *P. compressa* were found with significant break-points (*t-statistic <-2 or >2) between the control and the highest temperatures (30°C and 35°C). The red line is the median of the bootstrapped regression fit at each temperature, and shaded red areas indicate the bootstrap confidence intervals around the segmented regression fit. B) Network plots for one example cluster (cluster 9, *P. compressa*) showing the changes in hub genes identity at the control temperature and high (30°C and 35°C) temperature treatments. Each colored dot represents a hub gene at a specific temperature, which may stay a hub gene at a different temperature (same color within the network) or may be replaced by a new gene (different color within the network). See Fig. S18 for the hub rearrangement patterns across temperatures for all species.

Global network comparisons using the Jaccard index revealed no consistent pattern of overall network preservation or rewiring across temperature contrasts in any species (Fig. S17). However, examination of hub genes identity uncovered major temperature-dependent turnover (Fig. 3B), revealing extensive reorganization of core regulatory architecture that is not captured by global network metrics. This wholesale replacement of network hub genes showed overall the same pattern in all three species, although the number of hub genes involved is fundamentally different (Fig. S18). Indeed, stress induces cellular networks rearrangement involving the establishment of different hub genes in model taxa (94, 95). This phenomenon has been previously observed in environmental and stress responses of other eukaryotes, but so far it has not been reported for corals (96–99).

Enrichment analysis of the biological processes and molecular functions within hub genes between the species with the least plasticity, *P. compressa* and *M. capitata* (categories in bold in Fig. 4 and Fig. S19), and the highest plasticity, *P. acuta*, identified enrichment for ribosome biogenesis, a hallmark for coral response to temperature (33, 100–102), wound healing, amino acid metabolic process, essential for stress tolerance (103–106), and maintenance of cell polarity, which plays a critical role during energetic cellular stress (107–109)(Fig. S19) in *P. compressa* and *M. capitata* at the control temperature. These are supportive of frontloaded mechanisms contributing to maintaining elevated thermal stress resistance in *P. compressa* and *M. capitata.* For these species, hub gene functions at high temperatures shift toward enrichment of other stress response functions such as inflammatory response and programmed cell death, but also to energy generation functions such as carbohydrate and lipids metabolic processes (Fig. 4), reinforcing the importance of coral energetic resources, of which *P. compressa* and *M. capitata* show high levels (Fig. S6), in supporting vital processes under stress (75). Conversely, the functional enrichment landscape of *P. acuta* reveals multiple stress related functions with significant enrichment already at 30°C, such as wound healing, DNA repair, ribosome biogenesis, programmed cell death, immunity, amino acid metabolic processes and detoxification (Fig. 4). The latter, highlights the cost of the coral host to neutralize the negative effects of free radicals overproduced by the symbiotic algae during thermal stress, which are a major cause of coral bleaching (110–112). One of the top hub genes at 35°C for *P. acuta* is indeed an oxidative stress response gene (dihydropteridine reductase-like protein, (113); Table S9), indicating that *P. acuta* has started to face the initial stages of bleaching before the other two species, which is also the case in long term and field bleaching assessments (81, 114). These findings reveal that coral holobionts may respond differently to future changing climate, and that thick-tissued corals (e.g., *P. compressa* and *M. capitata*) will likely have an advantage because of their higher level of energy reserves (14), and as we reveal here, the internal conditioning of their molecular response mechanisms.

**Fig. 4.**
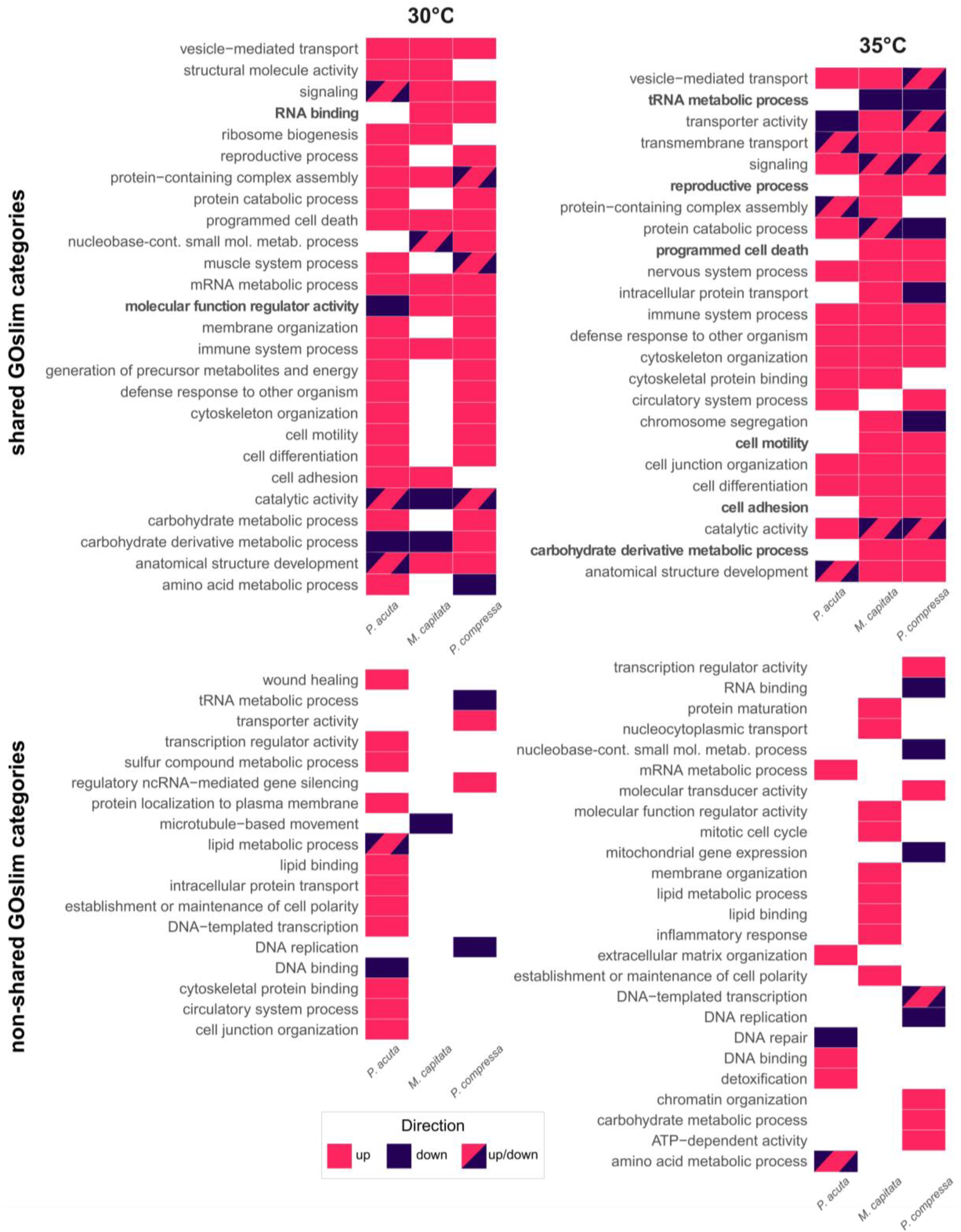
Biological processes and molecular functions enriched at high temperatures. Shared and non-shared (species-specific) GO slim categories enriched among hub genes at 30°C (heatmaps on the left) and 35°C (heatmaps on the right) from significant WGCNA clusters/modules (clusters/modules within the same species are grouped together). Colors indicate the predominant direction of eigengene expression changes at each temperature: positive module-trait correlation (up, pink; the majority of clusters/modules for that species show positive correlation), negative module-trait correlation (down, purple; the majority of clusters/modules for that species show negative correlation), mixed change (up/down, pink/purple; half the clusters/modules for that species show positive correlation and the other half shows negative correlation). GO slim categories in bold are shared between *M. capitata* and *P. compressa*, but not *P. acuta*.

### Isoform switching influences gene network architecture by establishing new hub genes

We further explored the mechanisms underlying the hub genes rearrangement at high temperatures by analyzing genome-wide isoform switching patterns. The percentage of the genome undergoing isoform switching at the combined temperature comparisons (control versus 30°C, control versus 35°C, 30°C versus 35°C) greatly varied across species, corresponding to 12.25% for *P. acuta,* 4.30% for *M. capitata* and 1.90% for *P. compressa.* This shows that *P. acuta* exhibited a much higher proportion of functional isoform switches as compared to the other two species, mirroring *P. acuta*’s larger transcriptome shift. Enrichment of alternative splicing events also differed across species (Fig. 5A), and we observed that *P. acuta* exhibits a lower threshold temperature for enrichment of AS events in switching isoforms, as well as the overall largest enrichment of AS events at high temperatures. Intron retention (IR) is among the most enriched events in the isoforms that switch at both 30°C and 35°C in *P. acuta* (Fig. 5A). In humans, mice and plants, IR is commonly associated with post-transcriptional regulation of stress and immune responses (115–117). This AS event increased in plants subjected to heat stress that were not subjected to a prior heat stress (stress-primed), whereas plants that were heat stress-primed produced similar splicing patterns to those of control plants (118), which is the case here for *P. compressa* that does not show enrichment of AS events in the switching isoforms at high temperatures (Fig. 5A). This supports our hypothesis of coral thermopriming based on life-history traits such as tick tissues and perforate skeletons (14, 24, 73). Undoubtedly, AS events are central to tune organismal responses to environmental differences, and the changes in isoform diversity through AS have been increasingly recognized as key players in phenotypic plasticity in a variety of organisms (119). Here, we provide the first evidence that corals exhibit species-specific patterns of alternative splicing events and isoform usage in response to temperature stress, adding a new dimension to our understanding of splicing in coral environmental response.

**Fig. 5.**
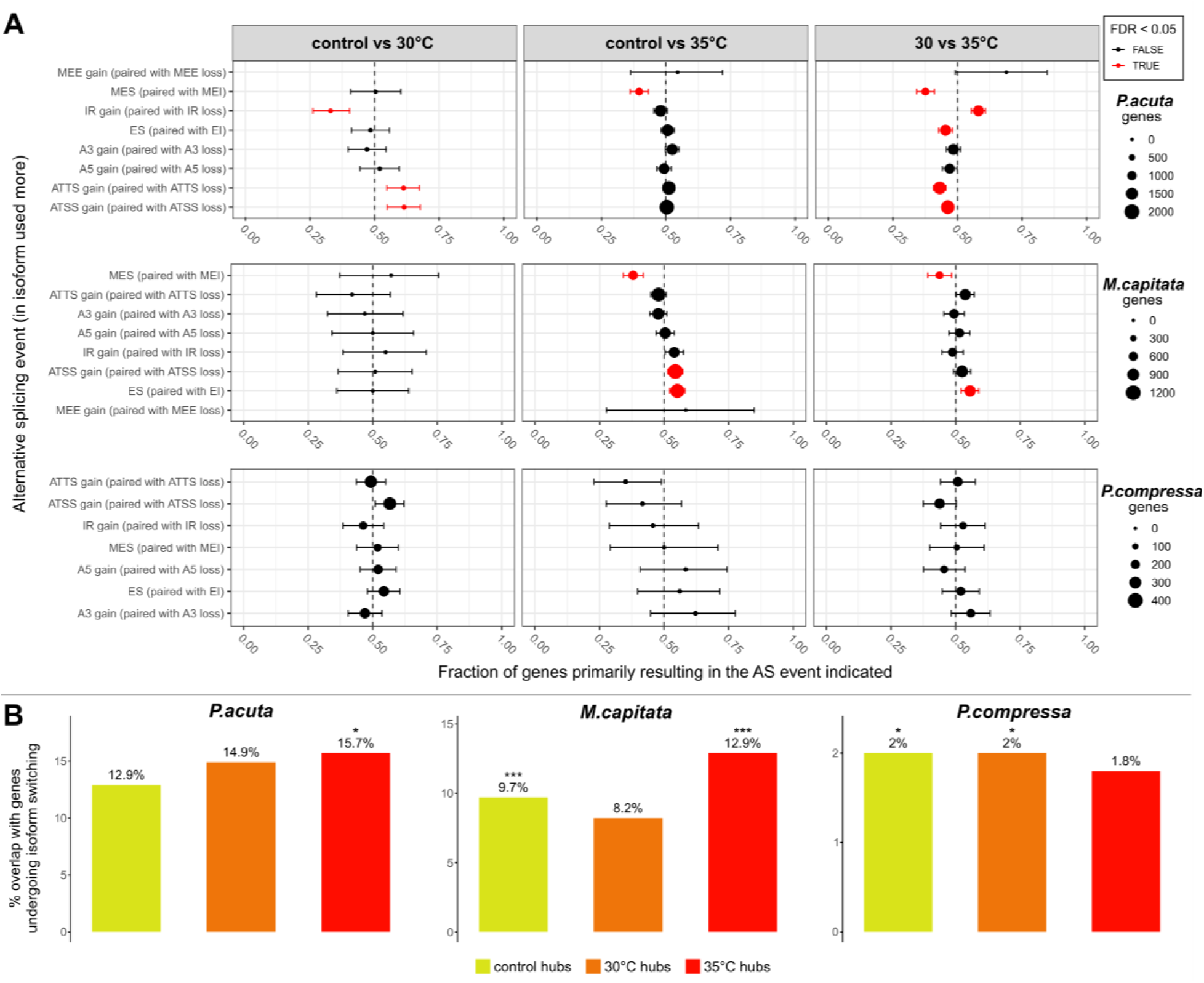
Enrichment of alternative splicing events in genes undergoing isoform switching and hub genes involvement in switching. (A) Alternative splicing events enrichment involved in isoform switches. The x-axis shows the fraction of genes with enrichment of a specific alternative splicing event, with opposite events shown together. Bars represent 95% confidence intervals, dot size represents the number of genes involved in switching events (different scales across species as each species was analyzed separately). Red dots represent events that are statistically significant, i.e. false discovery rate (FDR) < 0.05. For *P. compressa*, the lack of enrichment of alternative splicing events means that individual splicing events are not significantly enriched in genes undergoing isoform switching as compared to all isoforms (not just switching) in all temperature comparisons. IR intron retention, A5 alternative 5′ donor site (changes in the 5′end of the upstream exon), A3 alternative 3′ acceptor site (changes in the 3′end of the downstream exon), ATSS alternative transcription start site, ATTS alternative transcription termination site, ES exon skipping, EI exon inclusion, MES multiple exon skipping (skipping of > 1 consecutive exons), MEI multiple exon inclusion, MEE mutually exclusive exons. (B) Percentage of hub genes that undergo switching. Numbers above each bar represent the percentage overlap between unique hub genes at each temperature and genes undergoing isoform switching at high temperature. Asterisks indicate statistical significance of the overlap (*p < 0.05, **p < 0.01, ***p < 0.001; see Table S10 for statistical tests details).

Integration of isoform and network analyses demonstrated that hub genes turnover at high temperature is tightly linked to species-specific patterns of isoform switching. Across species, new hub genes at high temperature either arise from genes that switched isoforms or from genes that remained isoform-stable (Fig. 5B and Table S10), revealing distinct modes of network leadership reorganization. Similar associations between patterns of spliced isoforms and changes in hub genes identity have been reported for humans, found to be associated with functions dependent on tissue type (61) and, recently, with disease (62). The distinctly different patterns in the overlap between hub genes and genes that switch isoform at high temperatures found here (Fig. 5B and Table S10) reveal species-specific thermal response mechanisms. A stress-induced reorganization response type is prominent in *P. acuta*, where control hubs don’t switch, indicating that these network leaders remain stable, and 35°C hubs emerge from switched isoforms, indicative of stress-responsive reorganization of network leadership. This could indicate either reactive-only stress sensitivity, or specialization of a dedicated stress machinery. A multilayered response type was present in *M. capitata*, with control hubs switching isoform when temperature increases, and 35°C hubs emerging from genes that underwent isoform switching at lower temperatures and became new hubs. This species shows a flexible, dynamic response, with isoform plasticity being at the core of the gene network rewiring across temperatures. And finally, a continuative modulation response type occurs in *P. compressa*, with control and 30°C hubs switching isoform at higher temperatures, and 35°C hubs coming from isoforms that didn’t switch. Here, genes that were hubs at control and 30°C temperature change their isoform usage when temperature increases, and network rewiring at 35°C is determined by constitutively stable genes (that do not change isoform usage). This suggests that *P. compressa* utilizes a continuous isoform adjustment across temperatures while fine-tuning thermal response through dedicated hub genes. Overall, these findings reveal that isoform switching serves dual roles in the coral response to temperature changes: functional acclimatization, by changing protein properties under cellular stress (120), and network reorganization, reshuffling gene regulatory hierarchies. Moreover, the hub genes patterns reveal that coral species use fundamentally different network topology strategies under thermal stress, with species-specific differences reflecting divergent thermal niches.

### Integrative view of coral thermal tolerance and implications for reef future

Taken together, our findings reveal how coral thermotolerance is underpinned by intertwined physiological, gene regulatory, and post-transcriptional mechanisms in species-specific ways. *P. compressa* appears to rely on high basal energy reserves, gene frontloading and isoform switching of pre-existing hub genes, reflecting a strategy of robustness that is supported by the symbiont photosynthetic performance at high temperatures. *P. acuta*, in contrast, undergoes extensive transcriptomic shifts and isoform switching at high temperatures, indicating a reactive mode of thermal response but potentially costly strategy in the long term, consistent with the holobiont lower thermal threshold. *M. capitata* exhibits intermediate traits, relying on high basal energy reserves, gene frontloading and high isoform plasticity, but shows elevated photosynthetic sensitivity to high temperature, suggesting a mismatch between host and symbiont thermal tolerance.

These results provide a mechanistic framework to explain why closely co-occurring coral species exhibit different bleaching susceptibilities and survival trajectories under marine heatwaves. We have identified isoform switching as a previously unrecognized yet central element of coral thermal performance, reshaping gene regulatory hierarchies under stress and contributing to differential resilience among reef-building species. Future studies are needed to track cross-species AS and isoform switching patterns in corals across subsequent thermal stress exposures and recovery phases, to test if thermopriming based on life-history traits aids long-term survival. This would inform robust projections of shifts in coral community composition under continued climate warming, which are vital for policymakers to achieve effective strategies for coral reef management and restoration, helping us reach our climate and biodiversity goals (121).

## Methods

### Coral Collection

Coral colonies (n=8 per species named here as colonies A, B, C, D, E, F, G, H) of each of three coral species *Montipora capitata*, *Pocillopora acuta*, and *Porites compressa*, were collected from Reef 13 (1-3 meters) by snorkeling under a Special Activity Permit (SAP 2024-24, Division of Aquatic Resources, Honolulu, HI) on the 19^th^ of September 2023 using a hammer and chisel. Colonies were transported to the Hawaiʻi Institute of Marine Biology (HIMB) and held in a tank with sand filtered seawater (Pentair Triton II TR100 sand filter) under a natural light cycle with shade cloth with an average tank PAR of 493 µmol m^-2^ s^-1^. Corals were cut into 10 replicate fragments using bone cutters. Fragments were labeled according to genotype and fragment number (e.g., A1-A10, B1-B10, …), affixed to aragonite fragment plugs using cyanoacrylate glue (IC-Gel Insta Cure Cyanoacrylate Coral Frag Glue, Bob Smith Industries), and left to acclimate until their day of the experimental measurements (minimum 2 days, 21^st^ of September 2023, to a maximum of 10 days, 29^th^ of September 2023).

### Experimental setup

Following acclimation, each fragment was placed into individual acrylic respiration chambers (∼365 mL) with a magnetic stir bar and seawater. Each photosynthesis-irradiance (PI) curve run included coral fragments (n = 8 to 20 per run) and empty chambers with seawater as blanks (n = 2 per run). Chambers were placed in temperature-controlled tanks with a recirculating pump (Hydor Pico 300), chiller (SEAAN Aquarium, B08MXWLD59) and heaters (Aqueon Pro Heater, AP300W-H). Chambers were mixed with stir bars (speed of 310 rpm) on magnetic stir plates and held in a water bath maintained at the desired temperature via the above heaters and chillers plugged into an Apex Aquacontroller and EB32 (Neptune Systems). In order to measure oxygen flux corrected for temperature, a fiber-optic oxygen probe (PreSens dipping probe; DP-PSt7-10-L2.5-ST10-YOP) and temperature probe (PreSens Pt1000) were inserted into each chamber. Chambers were exposed to 10 increasing light levels for 10 minutes each (0 [dark], 34, 121, 315, 492, 638, 948, 1048, 1192, and 1378 µmol photons m⁻^2^ s⁻^1^; PAR) with LED aquarium lighting (Prime 16HD Reef Aquarium Lights, AquaIllumination) as quantified by an underwater cosine corrected sensor (MQ-510 Quantum Meter, spectral range of 389-692 ± 5 nm, Apogee Instruments, Logan, UT, USA) at each chamber position prior to each day of measurements. Oxygen concentration (µmol L^-1^) and temperature (°C) were logged every 1 sec.

While in the chambers, coral were simultaneously exposed to 6 temperature categories (12, 18, 25, 26.8, 30, 35) with 26.8 °C being the control temperature, i.e. the mean ambient temperature measured at this time of the year (August through October) in Kāne‘ohe Bay (measurements from the Pacific Islands Ocean Observing System, PacIOOS, averaged for the months of August to October 2022). For each of the three species, n=8 coral fragments were measured as replicates for each temperature category, with each replicate coming from a different parent colony (putative genotypes A to H; Figure S20). Each coral sample was only exposed to a single temperature, and the order of the temperature was randomly selected across the 11 PI curve runs needed to complete the total of 134 coral samples. The measurements were made between the 21st and the 29th of September 2023. Actual temperatures recorded in the chambers during each run are reported in Table S11. After PI curve runs were completed, fragments were immediately placed in sterile whirl pak bags, snap-frozen in liquid nitrogen, transported into a dry shipper to the University of Rhode Island, and stored at -80°C until processing for physiological and molecular analyses as described below (see Fig. S20 for experimental design overview).

### PI and TPC curves

Rates of oxygen flux were calculated for each light level interval using localized linear regressions (alpha=0.2, percentile rank method) in the ‘LoLinR’ package (122) from oxygen measurements at 20 sec intervals (to reduce noise in the data and to allow for processing of local linear regressions through the large dataset) and then normalized to chamber volume to produce µmol O_2_ sec^-1^. Mean oxygen evolution rates were calculated for blank chambers for each light level and run and subtracted from coral oxygen evolution rates to normalize for oxygen flux in the seawater. Rates were then normalized to each fragment surface area for units of µmol O_2_ cm^-2^ s^-1^, using surface area measurements obtained using the wax dipping method (123)(see below). Oxygen evolution rates (µmol O_2_ cm^-2^ s^-1^) from each light level interval were used to generate PI curves and a quadratic equation (Equation 1; (124)) was used to identify PI curve characteristics of each species at each experimental temperature.

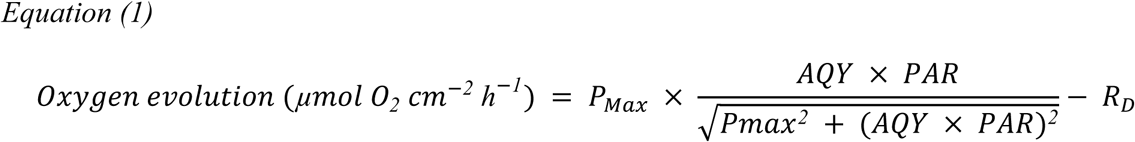

Where *P_Max_* is the maximal photosynthetic rate, *AQY* is the apparent quantum yield, *PAR* is the light value, and *R_D_* (µmol O_2_ cm^-2^ h^-1^) is dark respiration. In addition, saturating irradiance *I_K_* (light value (PAR) at which initial slope crosses *P_Max_*) was calculated as in Equation 2.

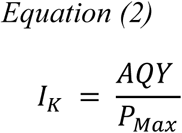

PI curve parameters *P_Max_*, AQY, *I_K_*, and *R_D_* were extracted for each colony at each of the 6 temperatures. Additionally, hourly *P_Max_*:*R_D_* ratios were calculated for each species and plotted across temperatures.

Individual thermal performance curves for *P_Max_* were fitted to the Pawar model (125), a modified version of the Sharpe-Schoolfield high temperature inactivation model (126, 127) that explicitly models the optimum temperature using the ‘rTPC’ package (128) with the following Equation 3:

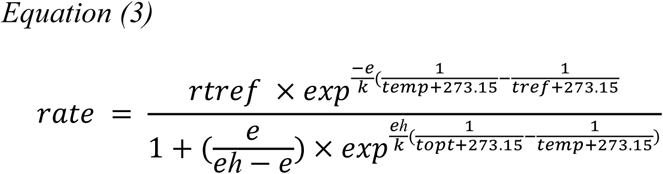

where *k* is Boltzmann’s constant (8.62e-05), *e* is the activation energy (eV), indicative of the steepness of the slope leading up to the thermal optima, *temp* is temperature in degrees centigrade (°C), *eh* is the deactivation energy (eV) which characterizes temperature-induced decrease in rates above the optimum temperature *topt* and *rtref* is the rate at the standardized temperature, where no low or high temperature inactivation is experienced. Fits were determined using the ‘nls_multstart’ function in the ‘nls.multstart’ package in R statistical software (v3.2.0)(129), which generates multiple start values and fits many iterations of the model using the Levenberg-Marquardt algorithm implemented in ‘nlsLM’. Measurement of uncertainty for the curve fit was done using the ‘weights’ argument within ‘nls_multstart’, to perform a weighted non-linear least squares regression as implemented in the rTPC package. To obtain confidence intervals around the curve fit prediction, the model was re-fit using the function ‘minpack.lm::nlsLM()’, with the coefficients of ‘nls_multstart()’ as the start values and using the weights as a correction to the Pearson residuals of the original model fit. The ‘Boot()’ function from the R package ‘car’ (130) was then used to resample the data and re-fit the model 999 times, generating 95% confidence intervals around both the fit prediction and the derived model metrics.

The work by (127) originally suggested using a *tref* of 25°C, considered appropriate for most poikilotherm organisms. Given that the *tref* should be well below the peak of the rate of interest (131), we chose 18° as *tref* for the model fit of *P_Max_*, by selecting the closest temperature to 25°C before the rate performance peak for all 3 species (Fig. S21). For *R_D_*, Fig. S21 shows that the rate for this PI-derived parameter does not possess the shape typical of a TPC with a well-defined peak in all 3 species. Therefore, a different model within the ‘rTPC’ package was used, namely the ‘sharpeschoollow_1981’ model (with *tref* of 25°C), a modified version of the classical Sharpe-Schoolfield model that only accounts for low-temperature inactivation (*el*). This is because in our *R_D_* data there is no thermal optimum but only data points before the TPC peak, so any estimate of metrics related to high-temperature would be imprecise (131)(see scripts in the github repository for more details).

To summarize multiple TPC metrics, we standardized each metric estimate across species using Equation 4:

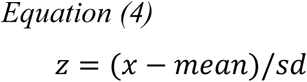

where z is the standardized metric, *x* is the metric for a given species, *mean* is the mean across all species, and *sd* is the standard deviation of that metric across all species. For metrics where higher values indicate greater thermal tolerance (e.g., *ctmax, topt, breadth, rmax, rtref*)(65, 127, 131–135), standardized values were used as-is; for metric where lower values indicate greater tolerance (e.g., *e, eh, el, ctmin*)(127, 134, 135), the sign of the standardized value was reversed. To visualize and compare the multivariate thermal tolerance profiles of each species, we used radar plots generated using the ‘fmsb’ R package. For each species, we plotted the standardized and directionally oriented values of all thermal performance metrics included in the composite score calculation. Metrics where higher values indicate greater thermal tolerance were plotted as-is, while those where lower values indicate greater tolerance were sign-flipped, ensuring that higher values on all axes consistently represent greater thermal tolerance. This approach allows for side-by-side comparison of species’ relative performance across multiple metrics, with each axis representing a standardized metric and each species’ profile shown as a colored polygon. Radar plots were generated using the ‘fmsb’ R package.

To visualize multivariate differences in TPC metrics for *P_Max_* among coral species, we performed a principal component analysis (PCA) using the ‘prcomp’ function in the ‘stats’ package. For each metric, we generated 1000 bootstrap samples by drawing from a normal distribution defined by the metric’s mean and confidence interval, to capture uncertainty in metric computation. All TPC metrics were standardized (z-scored) prior to PCA to ensure equal weighting. The PCA was performed on the combined bootstrapped dataset, and the resulting scores were plotted as a cloud for each species in PCA space. Variable loadings (arrows) were overlaid to indicate the direction and strength of each TPC metric’s contribution to the principal components.

### Physiology sample processing

Coral tissue was removed from the skeleton of each fragment into clean ziplock bags using an airbrush (Iwata Eclipse HP-BCS) filled with ice-cold 1X Phosphate Buffer Saline (PBS). The resulting tissue slurry was quantified for volume, placed into a 50 mL falcon tube and homogenized for 60 seconds (PRO Scientific Bio-Gen PRO200 Homogenizer). The tissue slurry was separated into 1.5 mL aliquots that were frozen at -20°C for chlorophyll and cell counts and at -80°C for protein and carbohydrates analyses. The remainder was frozen for tissue biomass measurements. The airbrushed coral skeletons were bleached and placed in a drying oven (Fisher Scientific Isotemp Oven) at 60°C for 24 hours before surface area measurements were taken as follows: pre-weighed wooden dowels of known dimensions were used to determine a standard curve of mass change of wax-dipped dowels against geometrically calculated surface area, according to the single wax dipping technique of (123). Surface area of bleached and dried coral skeleton was measured according to (123), by weighting the skeleton before and after wax dipping and using the standard curve from the wooden dowels. Assignment of samples to each physiological analyses was done as follows: protein, carbohydrate concentration and tissue biomass analyses were carried out only for control temperature samples (n=6-8 fragments per species and analysis) to assess baseline cross-species differences; chlorophyll concentration and algae density were carried out for all temperatures samples (n=6-8 fragments per temperature per species per analysis), to assess cross-species and within-species differences across temperatures. For each physiological analyses, a total of 8 genotypes were included.

### Total host protein

To quantify host total (soluble and insoluble) protein concentration, the 1.5 mL tissue homogenate aliquot was thawed on ice, vortexed to mix, 650 µL were pipetted into a new tube and centrifuged at 4°C at 13000 rpm for 4 minutes to separate the host from the algae. An aliquot of 500 µL of the supernatant was pipetted into a new tube and 10 µL of 1M NaOH were added, pH was confirmed to be ∼10 using pH paper. Homogenate samples were incubated at 50°C for 4 hours, with shaking at 300 rpm. After incubation, 0.1M HCl was added to the tube to neutralize the sample, until a pH of 7 was reached. 25µL of homogenate was added to each well on a 96 well clear flat bottom plate. Soluble protein content was measured using the Pierce BCA Protein Assay Kit (Thermo Fisher Scientific Cat #23227) with 25 µL of each sample or standard according to the manufacturer’s instructions. Protein content was calculated as absorbance at 562 nm against a bovine serum albumin (BSA) standard curve on a plate spectrophotometer (Synergy HTX Multi-Mode Reader, BioTek, USA). Values were normalized to homogenate volume and surface area for total protein concentration units of mg protein cm^-2^.

### Chlorophyll concentration

To quantify chlorophyll *a* and *c_2_* pigment concentration, a 1 mL tissue homogenate aliquot was thawed on ice, vortexed to mix, and centrifuged at 4°C at 5000 rpm for 4 min to separate the host from the algae. Supernatant was removed and replaced with 1 mL 1X PBS. This process (centrifugation and addition of PBS) was repeated 4 times. After the last centrifugation, supernatant was thoroughly removed and chlorophyll was extracted from each sample by adding 1 mL of 100% acetone to the symbiont pellets, vortexing, and then storing in a dark fridge at 4°C overnight. After the incubation, samples were centrifuged at 13000 rpm for 3 minutes and 200 μL of the extract was transferred to each of triplicate wells per sample in a 96-well quartz microplate (Hellma Analytics; Markham, Ontario Canada). The absorbance of each sample was measured at 630, 663, and 750 nm on a plate spectrophotometer (Synergy HTX Multi-Mode Reader, BioTek, USA). Chlorophyll *a* and *c_2_* concentrations were calculated from the equations for dinoflagellates in 100% acetone (Equation 5 and Equation 6 (136), and corrected for path length of the quartz plate (0.584 cm).

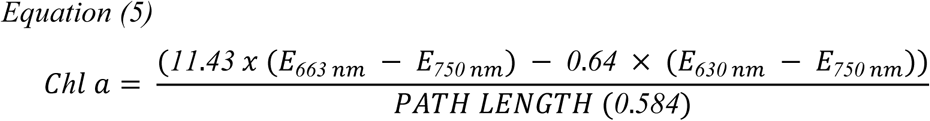

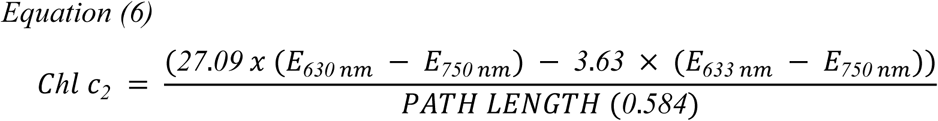

Chlorophyll *a* and chlorophyll *c_2_* content were normalized to skeletal surface area (µg pigment cm^-2^) and symbiont cell density (described below; µg pigment cell^-1^).

### Symbiodiniaceae density

To quantify symbiotic algal cell density, 0.650 mL tissue homogenate aliquots were thawed on ice, vortexed to mix, and centrifuged at 4°C at 5000 rpm for 4 min to separate the host from the algae. Supernatant was removed and replaced with 1 mL 1X PBS. This process (centrifugation and addition of PBS) was repeated 3 times. Algal cell numbers were determined following (137) using a Guava easyCyte HT flow cytometer (Millipore) with the InCyte v3.4 program (Millipore); algal cells were discriminated from host-cell debris using gates based on forward scatter and chlorophyll a fluorescence (detected after laser excitation at 488 nm). Three technical replicates were quantified for each sample in a 96-well plate, 240 µL of sample per well. Algal cells were also counted using a hemocytometer (n=5 samples per species), to ensure that the Guava was not underestimating algal cell numbers because of the decrease in the chlorophyll-a fluorescence per cell. The trendline generated for the Guava-hemocytometer counts shows an R-squared of 0.96 (Fig. S22). The concentrations of symbionts in each well were averaged across the replicates and normalized to homogenate volume and surface area to obtain cell cm^-2^.

### Host carbohydrates concentration

Host fractions for each sample were analyzed for total carbohydrates following the phenol-sulfuric acid method modified for use in 96-well plates (138), and 100 μL of each sample (host fractions of n = 22 samples, control temperature) were added to 900 μL of deionized water in 5 mL tubes. Carbohydrates were extracted by adding 25 μL of phenol and 2.5 mL of sulfuric acid to each sample or L-(-)-Glucose standards (Sigma-Aldrich cat. G5500) and allowed to incubate for 30 min. Following incubation, 200 μL of each sample and each standard were then added to 96-well plates in triplicate. Absorbance of each well was then read on a plate reader (Synergy HTX Multi-Mode Reader, BioTek, USA) at 485 nm. Carbohydrate concentration (mg mL^-1^) was determined against a standard curve of the L-(-)-Glucose standards as a function of absorbance at 485 nm and normalized to total host soluble protein (μg carbohydrate over μg host protein^-1^) and symbiont cell density (μg carbohydrate cell^-1^).

### Tissue biomass

Tissue biomass was quantified using the AFDW method. Specifically, 5 mL of tissue slurry was centrifuged at 13,000g for 3 min. A volume of 4 mL of the supernatant host tissue was removed and placed in pre-burned aluminum pan weighed on an analytical balance (Mettler Toledo, model ML203T/00). The samples were placed in a drying oven (Thermo Fisher Scientific, Heratherm General Protocol Oven, catalog no. 51028112) for 24 h at 60°C, weighed, and then placed in a muffle furnace for 4 h at 450°C (Thermo Fisher Scientific, Lindberg Blue M Muffle Furnace, catalog no. BF51728C-1). AFDW (mg cm^−2^) of the host fraction was calculated as the post-drying oven mass (dry mass)-post-muffle furnace mass, and final values were standardized to the airbrushed homogenate volume and normalized to surface area.

### RNA extraction, sequencing and reads processing

RNA was extracted from 3-4 samples per species and temperature (4 genotypes included in total) using the Zymo Quick-DNA/RNA Miniprep Plus (Zymo Cat# D7003, Zymo Research Corporation) kit following the manufacturer’s instructions. RNA quantity and quality were assessed with a Qubit 3.0 Fluorometer (Thermo Fisher Scientific) and by agarose gel electrophoresis. Total RNA samples were then sent to the Oklahoma Medical Research Foundation NGS Core (Oklahoma City, Oklahoma, USA) for library preparation and sequencing. cDNA libraries were constructed using polyA enrichment with the Watchmaker Genomics RNA Library Prep Kit and were sequenced on an Illumina NovaSeq X Plus in a 2x150bp Paired End (PE) configuration, targeting 40 million reads per sample (20 million reads in each direction).

Raw reads were trimmed and filtered using fastp (v0.23.2)(139) to remove low-quality reads and adapters. Cleaned reads were aligned to the *P. acuta*, *M. capitata* and *P. compressa* host genome assemblies (140) using STAR (v2.7.11b)(141) in twoPass Basic mode and with filtering flags --outFilterMismatchNmax 10 --outFilterMismatchNoverLmax 0.3 –outFilterScoreMinOverLread 0.66 --outFilterMatchNmin 0. Samples with <5 million mapped reads, that is a typical read depth threshold to obtain reliable statistical power in differential expression analysis (142), were excluded from downstream analysis. This resulted in the exclusion of one sample from the 18°C treatment and one sample from the 26.8°C treatment for *P. acuta*, and of one sample from the 35°C treatment for *P. compressa.* Additionally, cleaned reads were mapped with STAR, using the same options as the host, to concatenated *Cladocopium* spp. genomes (143–146) for *P. acuta* and *P. compressa* and to concatenated *Cladocopium* spp. and *Durusdinium* spp. genomes (147, 148) for *M. capitata*, according to the Symbiodiniacea populations found in these coral species in Hawai’i (149). Alignment rates to symbiont genomes were below 5 million reads for all samples of all species, therefore no further data analysis was carried out for symbiont-related gene expression.

Host aligned reads were assembled using StringTie (v2.2.1)(150) and GFFcompare (v0.12.6)(151) was used to assess the precision of mapping by comparing merged mapped GTFs to the *P. acuta*, *P. compressa* and *M. capitata* reference assemblies. Finally, gene count matrices for the three species were generated using the StringTie python script prepDE. To provide gene ontology (GO) annotation for enrichment analysis, functional annotation of *P. acuta*, *P. compressa* and *M. capitata* protein sequences (140) was performed by finding homologous sequences using the Diamond blastp function (v2.0.11)(152) against the NCBI nr database (version of 2024-02-07). Homologous sequences were mapped to UniProtKB and UniParc databases using the Uniprot mapping service (https://www.uniprot.org/id-mapping/) (153) to retrieve GO information. Additionally, homologous sequences were annotated using eggNOG (154), and the outputs of the Uniprot and eggNOG searches were merged to generate a compiled annotation.

### Genome-wide gene expression plasticity across species

Genes lengths for each species were extracted from the stringtie-generated GTF files using the R package GenomicFeatures (v1.54.4). Transcripts per million (TPMs) were computed using the stringtie-generated raw count matrix of each species and were normalized using gene length information. TPMs were then log transformed and filtered to remove low counts genes by retaining genes with TPM > 1. The ‘normalizeBetweenArrays()’ function from the R package limma was then used to further adjust for distribution differences between samples within each species. To enable cross-species comparisons, the program Broccoli (v1.3)(155) was used to infer orthologous pairs of proteins across the three species using default settings.

To compare gene expression plasticity in response to high temperature among species (33), a discriminant analysis of principal components (DAPC) was carried out using the normalized and filtered TPM counts to compare expression of all the expressed genes at 26.8°, 30° and 35°C across the three species using the ‘adegenet’ package (v2.1.11)(156, 157). A discriminant function was set by defining each temperature per species as groups (Mcap_26.8, Mcap_30, Mcap_35, Pacu_26.8, etc.). DAPC finds the linear combinations of genes (discriminant functions) that best distinguish these groups. To compare gene expression plasticity, the R package ‘MCMCglmm’ (v2.36)(158) was used to model DAPC scores as a function of high temperature plus the species-specific effect of being at higher temperature. From the posterior distributions of the Markov Chain Monte Carlo (MCMC) model parameters, species-specific effects of high temperature exposure were extracted and pairwise differences in absolute plasticity effects between species were calculated. MCMC-based posterior probabilities were computed as the proportion of posterior samples where the absolute plasticity effect of one species exceeded another. These probabilities indicate the likelihood that one species exhibits greater transcriptomic plasticity than another, with values close to 1.0 (100% probability) representing strong evidence of a difference between species. The first linear discriminant (LD1) scores resulting from DAPC were visualized using density plots created with the R package ‘ggplot2’ (v3.5.1). To quantify and visualize the magnitude of transcriptomic plasticity for each species, mean LD1 scores were calculated for control and elevated temperature treatments within each species. Arrows were overlaid on the density plots to indicate the direction and magnitude of the shift in gene expression from control to elevated temperature conditions, with arrow length proportional to the plasticity response. The difference in mean LD1 scores (elevated temperature minus control) was added to the density plots in correspondence of each arrow to provide quantitative comparisons of plasticity between species.

### Within-species WGCNA and break-points estimation of modules expression profiles across temperatures

Within the R environment (v4.3.2)(159), the gene count matrix of each species was filtered independently to remove low-count genes using the ‘pOverA’ filter function of the ‘Genefilter’ package (v1.84.0). Specifically, genes with less than 10 counts in at least 3-4 samples (minimum number of replicates per species per temperature) out of 21-24 samples were excluded. Filtered counts were then normalized using the variance stabilizing transformation (vst) in ‘DESeq2’ (v1.26.0)(160) and a PCA was conducted using the ‘plotPCA’ function to visualize sample-to-sample distances. Normalized counts were used to perform Weighted Gene Co-expression Network Analysis (WGCNA)(161) to identify modules (groups of genes) with distinct expression profiles across temperatures within each species. First, an unrooted hierarchical tree was built using the R function ‘hClust’ with “average” method to check for outliers. The function ‘pickSoftThresholding’ of the ‘’WGCNA package (v1.73) was used to explore values of soft threshold from 1 to 30, to construct a topological overlap matrix similarity network and assess gene expression adjacency. A soft thresholding power of five for *P. acuta* and *M. capitata,* and of seven for *P. compressa* were chosen (based on the recommended scale-free topology fit index of 0.8; (161)) and were used to construct the topological overlap matrix similarity network with adjacency of type “signed”, to keep track of the sign (negative or positive) of the co-expression information. The ‘dynamicTreeCut’ function was used to identify modules from the topological overlap matrix similarity network with a minimum module size of 30, and resulting modules with > 80% eigengene similarity were merged for all species. The flashClust “average” method was used to cluster the expression modules by eigengene similarity, and module-trait correlation was assessed by determining the correlation between genes and temperature conditions (genes significance) and the correlation between modules eigengene and genes expression profiles (module membership). A heatmap of the module-trait correlation values was plotted per each species with the package ‘complexHeatmap’ and modules were divided into clusters with the complexHeatmap function ‘row_split’ to highlight changes in modules expression by temperature condition.

The expression profile in each cluster of modules was summarized by generating plots of mean eigengene expression value per each temperature condition. The variation in eigengene expression of each cluster was analyzed using the ‘segmented’ package (v2.1-4)(162), which uses regression models to detect break-points (change-points) in the relationship between a numeric predictor (here, temperature) and a response (eigengene value). Break-points significance was determined by both the t-statistic, which test whether the change in slope is significantly different from zero, computed by the segmented package (break-point significant if t-statistic <-2 or >2)(163) and by comparing the breakpoint model to a simpler linear regression model using Akaike Information Criterion (AIC), to test if the inclusion of a break-point significantly improves the fit (lower AIC indicates a better model). For clusters with significant break-points, bootstrap (n = 1000 re-samplings) was used to estimate the confidence intervals around the regression model.

To further characterize thermal thresholds at higher temperature than ambient, gene networks for clusters with significant break-points between 26.8°C and 35°C were analyzed to assess network stability or rewiring at high temperatures. Specifically, the Jaccard index was computed to quantify the similarity of global gene network structure between the control and the other temperature treatments. For each cluster, binary adjacency matrices were constructed by thresholding absolute Pearson correlation coefficients between gene pairs at a specified edge threshold (0.6). Edges present in either condition were identified, and the Jaccard index was computed as the ratio of the number of shared edges to the total number of unique edges in both networks. This metric ranges from 0 (no shared edges) to 1 (identical networks), providing a measure of network rewiring or preservation across temperatures. Additionally, hub genes in each cluster at 26.8°C, 30°C and 35°C were found based on intramodular connectivity (genes with connectivity ranked at top 10% within a network), measured through the sum of absolute correlations, i.e. ranking genes by their overall co-expression with all other genes in the cluster. Hub genes’ identity was then compared between ambient and high temperatures (26.8°C vs 30°C, 26.8°C vs 35°C and 30°C vs 35°C). The top hub gene for each temperature treatment and cluster modules was identified for each species and searched in the Diamond blastp output described above to identify the NCBI protein name. Genes corresponding to uncharacterized proteins in the blastp output were blasted again using the NCBI blast tool, and the top hit was selected (see Table S9 for the hub genes list).

### Functional enrichment of top hub-genes

Gene ontology (GO) enrichment analysis of the unique hub genes at 26.8°C, 30°C and 35°C (hub genes that are not shared between the 3 temperatures) for each cluster was performed, separately for each species, using the R package ‘ViSEAGO’ (v1.18.0)(164). Overrepresentation testing was performed using the R package ‘TopGO’ (165) runTest function, with algorithm = “classic” and statistic = “fisher”, and the set of all expressed genes (for each species separately) was used as background. GO terms were considered significant if they obtained a p-value of <0.01. For the resulting enriched GO terms, slim categories were obtained using the function ‘goSlim’ of the R package GSEABase (v1.66.0)(166), using the GOslim generic obo as reference database (v1.2)(167). The full lists of enriched GO terms and slim categories are available in the github repository https://github.com/fscucchia/HI_PhotoPhysio_TPC_geneExpr.

### Isoform switching analysis

Transcript-level count matrices generated by StringTie (with discovery of novel isoforms) were analyzed for isoform switching using the ‘IsoformSwitchAnalyzeR’ package (v2.4.0)(168) in R. To ensure proper normalization for both library size and transcript length differences, transcript lengths were first extracted from the merged GTF annotation using the ‘GenomicFeatures’ R package. Raw transcript counts were normalized for library size using ‘calcNormFactors()’ function of the ‘edgeR’ package (4.2.2), followed by conversion to FPKM values using the ‘rpkm()’ function with transcript-specific lengths. Prior to isoform switching analysis, expression values were filtered using the ‘preFilter()’ function of the IsoformSwitchAnalyzeR package to reduce noise from lowly expressed genes and isoforms. Filtering included exclusion of single-isoform genes, as they cannot exhibit isoform switching by definition, exclusion of genes with low overall expression (< 0.1 FPKM) to ensure reliable isoform fraction calculations, exclusion of individual isoforms with expression levels below 0.1 FPKM. Additionally, isoforms contributing less than 0.1% to their respective gene’s total expression were excluded (default for the IsoformSwitchAnalyzeR package).

Differential isoform usage testing was conducted using ‘isoformSwitchTestDEXSeq()’, which implements a DEXSeq-based statistical framework (169) with limma-derived effect size calculations to control false discovery rates. Significant isoform switches were identified using an FDR threshold of 0.05 and a minimum change in isoform fraction (dIF) of 0.01. Following DEXSeq, open reading frames (ORF) were annotated using the reference GTF file for known isoforms, followed by de novo ORF prediction using ‘analyzeORF()’ for novel transcripts discovered by StringTie. Nucleotide and amino acid sequences were extracted for alternative splicing analysis and functional consequence analysis, including coding potential prediction (CPC2) and protein domain annotation (Pfam). Functional consequences of isoform switches were assessed using ‘analyzeSwitchConsequences()’, examining changes in protein domains, ORF length, coding potential, and splicing patterns between temperature conditions. The isoform switching analysis was then integrated with the hub genes data from WGCNA to identify overlaps between temperature-specific network hub genes and genes undergoing isoform switching with higher temperature, testing for statistical enrichment using Fisher’s exact tests (p < 0.05).

### Statistical analyses

Host and symbiont physiological and PI parameters were tested for normality using the Shapiro-Wilk test (normality assumed if p > 0.05) and confirmed with quantile-quantile plots, and for homogeneity of variance using the Levene test (p > 0.05). For parameters that passed the Shapiro-Wilk and Levene tests, one-way ANOVA was used to test differences between species or between temperatures within the same species, followed by Tukey test for post-hoc comparisons. For parameters that didn’t pass the tests, the Kruskal-Wallis test was used to test differences between species or between temperatures within the same species, followed by the Dunn’s test for post-hoc comparisons. All pairwise post-hoc comparisons were considered significant if the post-hoc p < 0.05.

To assess whether differences between TPC metrics were statistically significant across species, pairwise comparisons of confidence intervals using a Z-score-based test was performed. For each metric, the Z-score was calculated for the absolute difference between two point metrics, using the pooled standard error derived from the width of their respective 95% confidence intervals. Since a 95% confidence interval corresponds to 1.96 standard deviations above and below the mean in a normal distribution, the standard error was approximated by dividing the confidence interval width by 3.92 (i.e., 2×1.962). The pooled standard error was then used to compute a Z-score for the difference between two metrics, and the Z-score was converted to a corresponding p-value using a two-tailed normal distribution. All statistical tests were performed in R (v4.4.2) and RStudio (v2024.12.0). Statistical tests results are presented in Supplementary Tables S1-S8.

## Acknowledgements

As guests, we recognize and give thanks for the land and water resources of the ‘āina and the traditional owners of the land, kānaka ‘ōiwi, both past and present, as well as future generations, on which this experimental work was conducted in the Kāne‘ohe Ahupua‘a on O‘ahu in the islands of Hawai‘i. We are grateful for the logistical support provided by the Coral Resilience Lab and the Hawai‘i Institute of Marine Biology. The sequencing was procured and managed via Genohub.com. We would like to thank the University of Rhode Island High Performance Computing for bioinformatics support. We would like to thank Dr. Susanne Menden-Deuer and Dr. Pierre Marrec at URI Graduate School of Oceanography for the use of their Guava instrument. This study was supported by the Heising-Simons Foundation International Ltd. The authors declare no conflict of interest.

## Author Contributions

H.M.P. designed the research; H.M.P. and P.H. conducted field work and experiment; F.S. conducted lab work; F.S. analyzed the data; H.M.P. acquired funding; F.S. and H.M.P. wrote the first draft; F.S, H.M.P. and P.H. edited the manuscript. All authors approve the final manuscript.

## Competing Interest Statement

The authors declare no competing interests.

## Notes

https://github.com/fscucchia/HI_PhotoPhysio_TPC_geneExpr

### Competing Interest Statement

The authors have declared no competing interest.

