## Supplementary Figures for "Contrasting temperature-induced gene network rewiring and isoform switching underlie relative thermal tolerance of coral species"

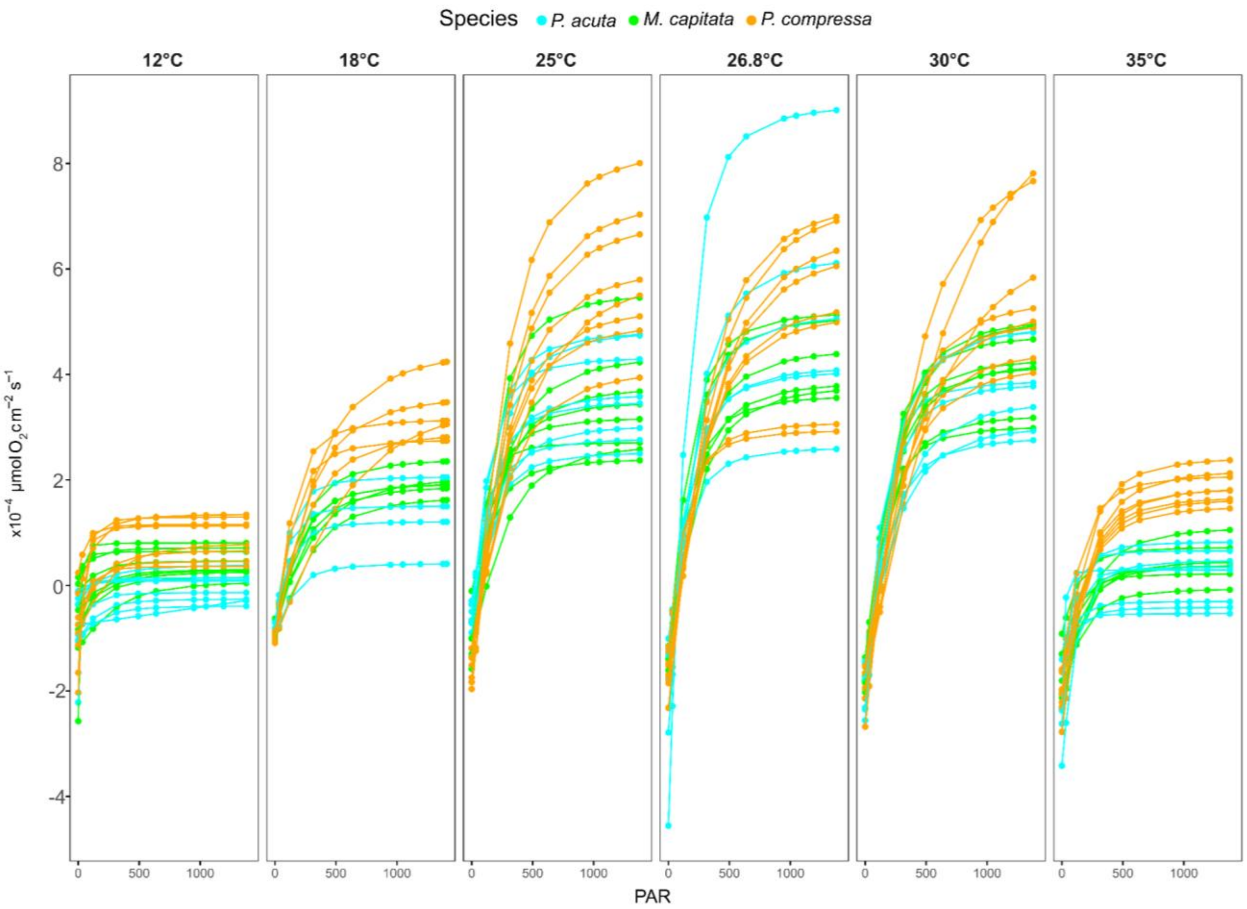


Figure S1. PI fitted curves across temperature treatments. PI curves showing photosynthetic oxygen evolution rates (μmol $O_2$ $cm^{-2}$ $s^{-1}$) as a function of photosynthetically active radiation (PAR, μmol photons $m^{-2}$ $s^{-1}$) for *M. capitata* (green), *P. acuta* (cyan), and *P. compressa* (orange) across the 6 temperature treatments (12°C, 18°C, 25°C, 26.8°C, 30°C, and 35°C). Each curve represents an individual sample, illustrating the range of photosynthetic responses within each species at each temperature.

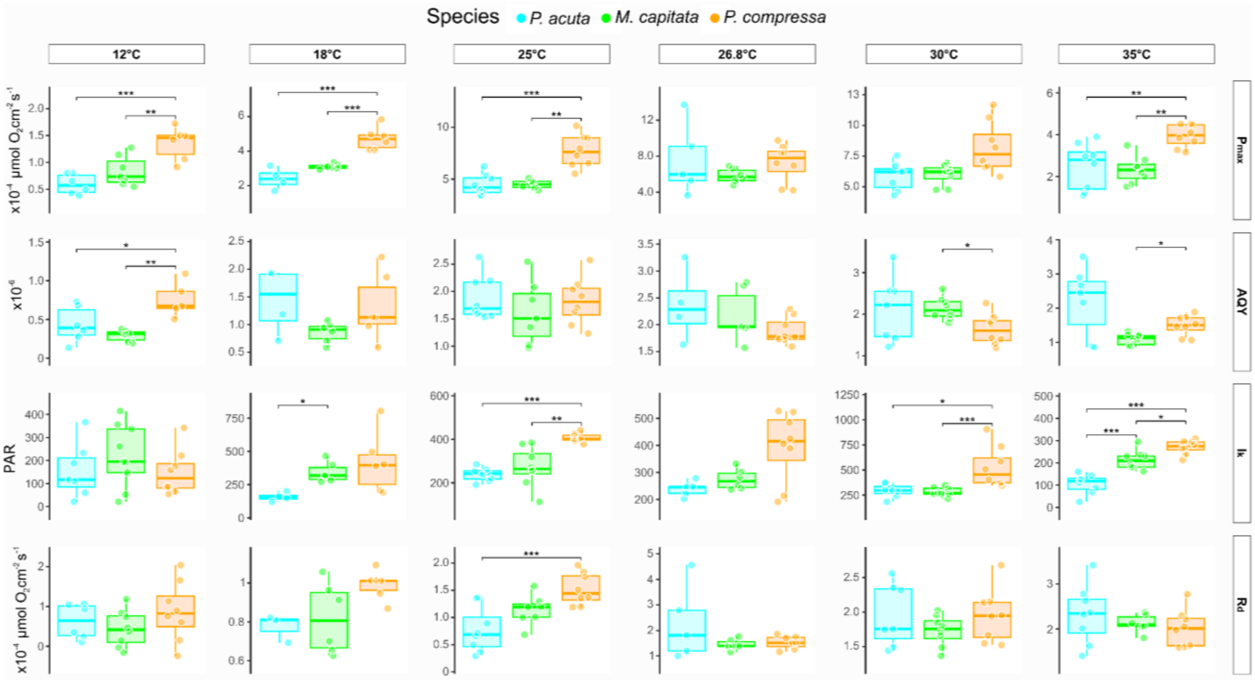


Figure S2. Interspecific comparison of PI-derived parameters across temperature treatments. Box plots comparing PI-derived photosynthetic parameters among *P. acuta* (cyan), *M. capitata* (green), and *P. compressa* (orange) at the 6 temperature treatments (12°C, 18°C, 25°C, 26.8°C, 30°C, and 35°C). Parameters shown are: Pmax (maximum photosynthetic rate, µmol $O_2$ $cm^{-2}$ $s^{-1}$), AQY (apparent quantum yield), Ik (light saturation parameter, PAR µmol photons $m^{-2}$ $s^{-1}$), and Rd (dark respiration rate, µmol $O_2$ $cm^{-2}$ $s^{-1}$). Each box represents the interquartile range with median values indicated by horizontal lines within boxes. Whiskers extend to the most extreme data points within 1.5 times the interquartile range, and individual data points (samples) are overlaid as circles. Horizontal brackets with asterisks indicate statistically significant differences between species pairs (*$p < 0.05$, **$p < 0.01$, ***$p < 0.001$; see Tables S1 and S2 for statistical tests details).

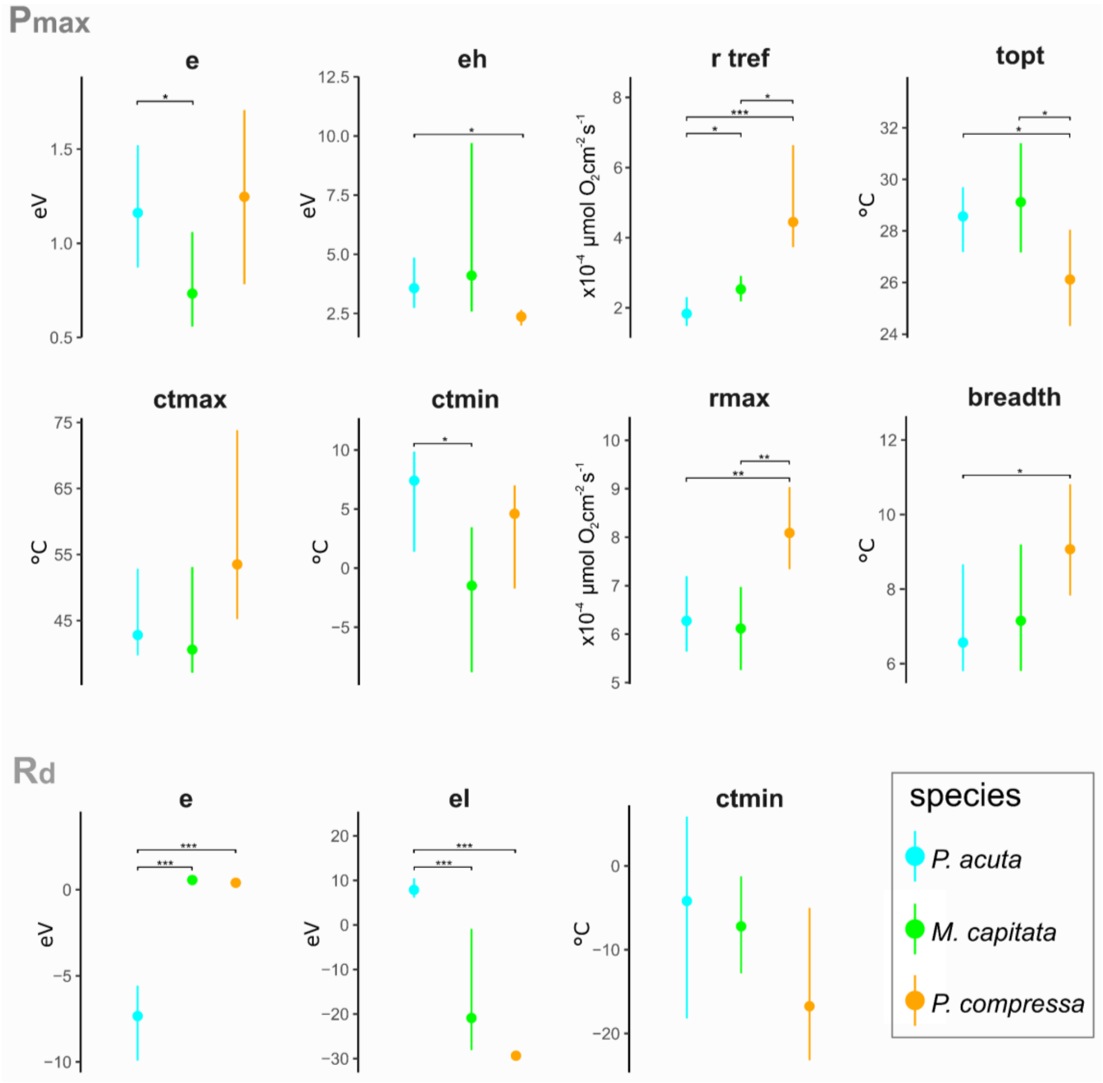


Figure S3. TPC metrics for maximum photosynthetic rate (Pmax) and dark respiration (Rd) across species. Mean values (±95% confidence intervals) of TPC metrics derived from temperature-dependent responses of Pmax and Rd for *M. capitata* (green), *P. acuta* (cyan), and *P. compressa* (orange). For Pmax: e (activation energy, eV), eh (deactivation energy, eV), r tref (rate at reference temperature, µmol $O_2$ $cm^{-2}$ $s^{-1}$), topt (optimal temperature, °C), ctmax (critical thermal maximum, °C), ctmin (critical thermal minimum, °C), rmax (maximum rate, µmol $O_2$ $cm^{-2}$ $s^{-1}$), and breadth (thermal performance breadth, °C). For Rd: e (activation energy, eV), el (low-temperature deactivation energy, eV), and ctmin (critical thermal minimum, °C). Horizontal brackets with asterisks indicate statistically significant differences between species pairs (*$p < 0.05$, **$p < 0.01$, ***$p < 0.001$; see Table S3 for statistical tests details).

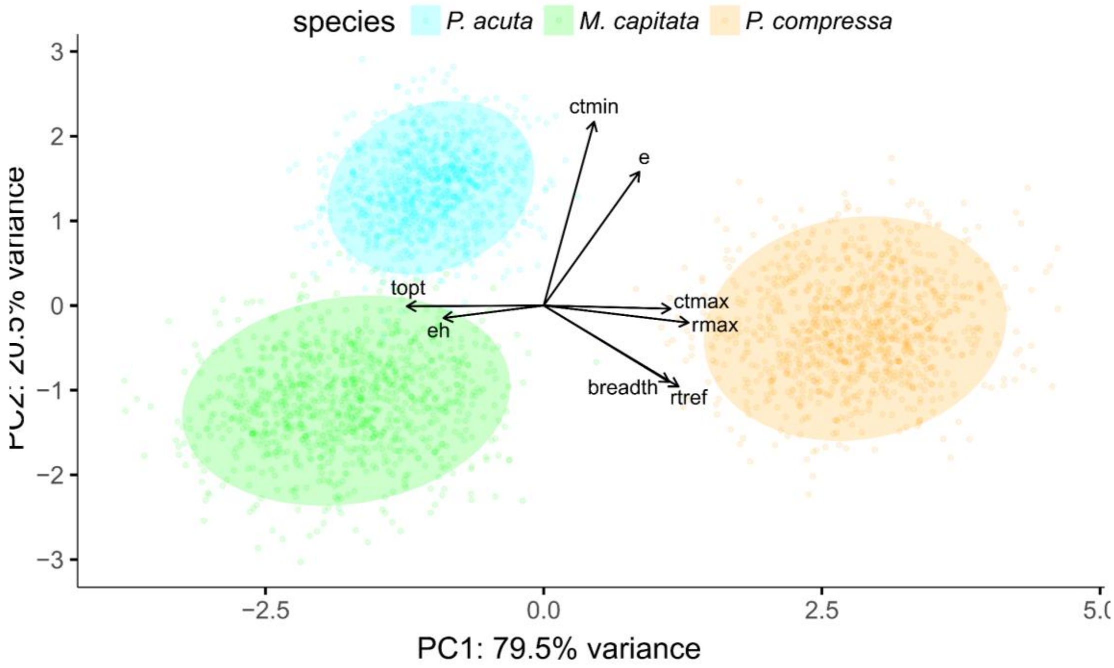


Figure S4. PCA biplot of bootstrapped TPC metrics for Pmax. Each coral species, *M. capitata* (green), *P. acuta* (cyan), and *P. compressa* (orange), is shown as colored point clouds, reflecting the uncertainty in metrics estimation. Each cloud represents the multivariate spread of TPC metrics for a given species, with ellipses indicating the 95% confidence region. Arrows indicate the loadings of individual TPC metrics on the first two principal components; the direction and length of each arrow show how strongly and in which direction each metric contributes to variation among species (loading on PC1/PC2). Each point: one bootstrap sample for a species (reflects parameter uncertainty).

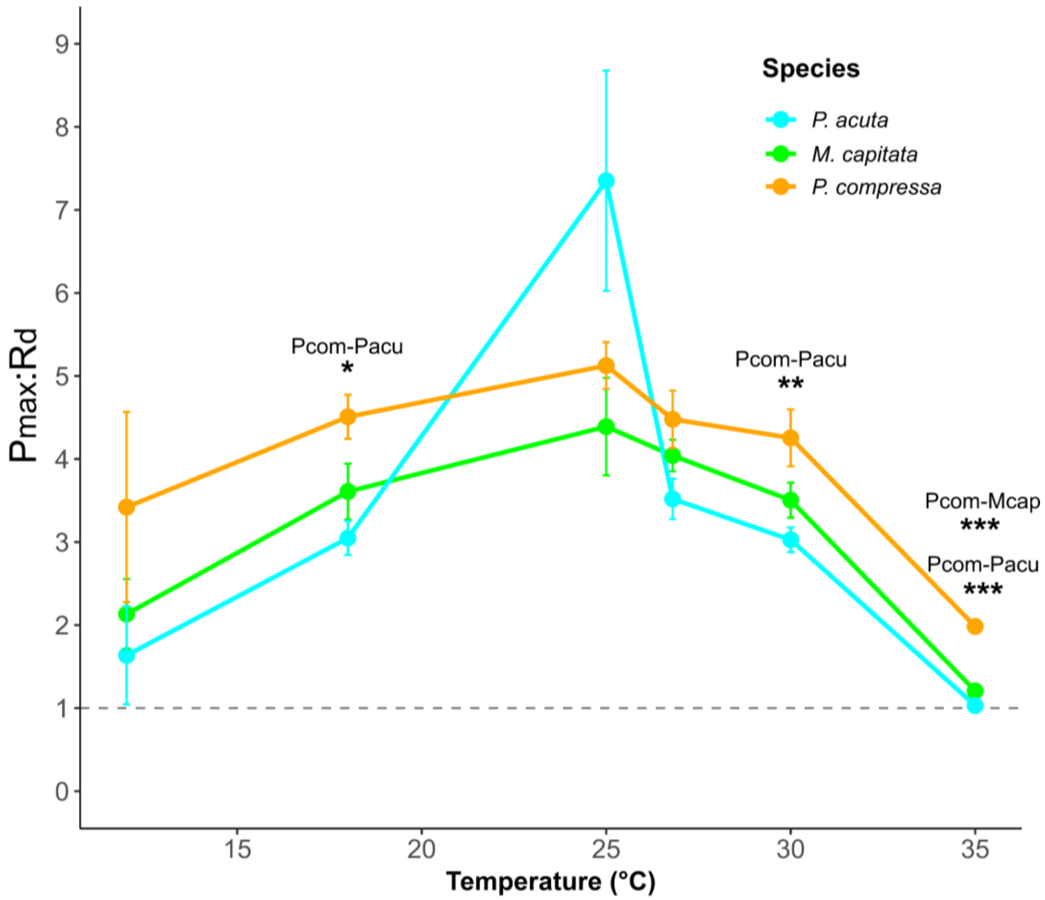

Figure S5. Hourly max gross photosynthesis to respiration ratios by temperature. For each species, Pmax:Rd ratios were calculated and displayed as mean values ± standard errors. Dashed horizontal line is at a Pmax:Rd = 1. Values < 1 indicates that the dark respiration rate is higher than the gross photosynthesis rate. Asterisks indicate statistically significant differences between species pairs at each temperature (*p < 0.05, **p < 0.01, **p < 0.001; see Tables S1 and S2 for statistical tests details).

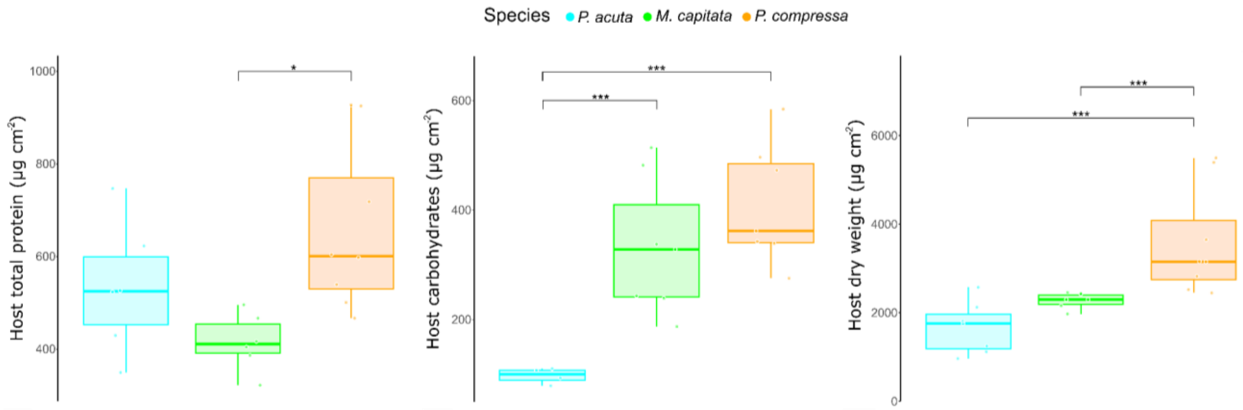


Figure S6. Coral host physiological parameters at control temperature (26.8°C). Box plots comparing host physiological metrics among *M. capitata* (green), *P. acuta* (cyan), and *P. compressa* (orange), measured at the control temperature of 26.8°C. Parameters include host total protein (µg cm$^{-2}$), host carbohydrates (µg cm$^{-2}$), and host dry weight (µg cm$^{-2}$). Each box represents the interquartile range with median values indicated by horizontal lines within boxes. Whiskers extend to the most extreme data points within 1.5 times the interquartile range, and individual data points (samples) are overlaid as circles. Horizontal brackets with asterisks indicate statistically significant differences between species (*$p < 0.05$, ***$p < 0.001$; see Tables S4 and S5 for statistical tests details).

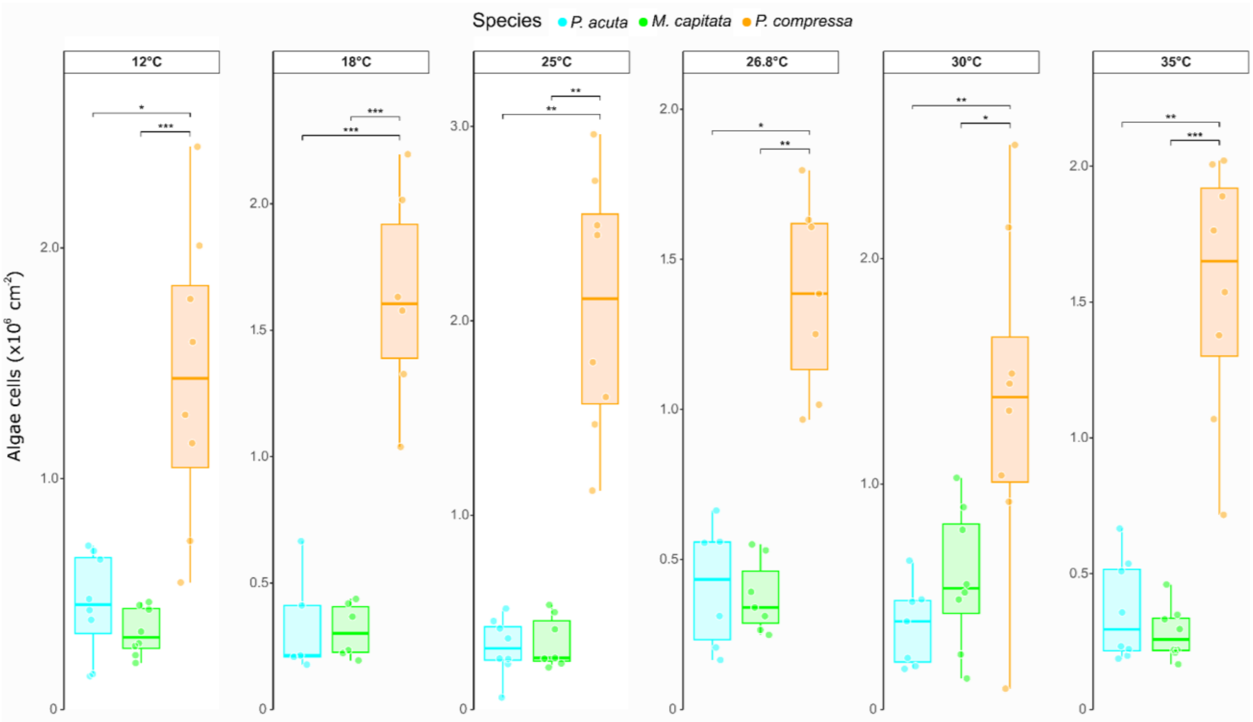


Figure S7. Cross-species algae density changes across all temperature treatments. Box plots showing algae cell density per skeleton surface area ($\times10^6$ cells $cm^{-2}$) for *M. capitata* (green), *P. acuta* (cyan), and *P. compressa* (orange) across the six temperature treatments (12°C, 18°C, 25°C, 26.8°C, 30°C, and 35°C). Each box represents the interquartile range with median values indicated by horizontal lines within boxes. Whiskers extend to the most extreme data points within 1.5 times the interquartile range, and individual data points (samples) are overlaid as circles. Horizontal brackets with asterisks indicate statistically significant differences between species pairs (*$p < 0.05$, **$p < 0.01$, ***$p < 0.001$; see Tables S6 and S7 for statistical tests details).

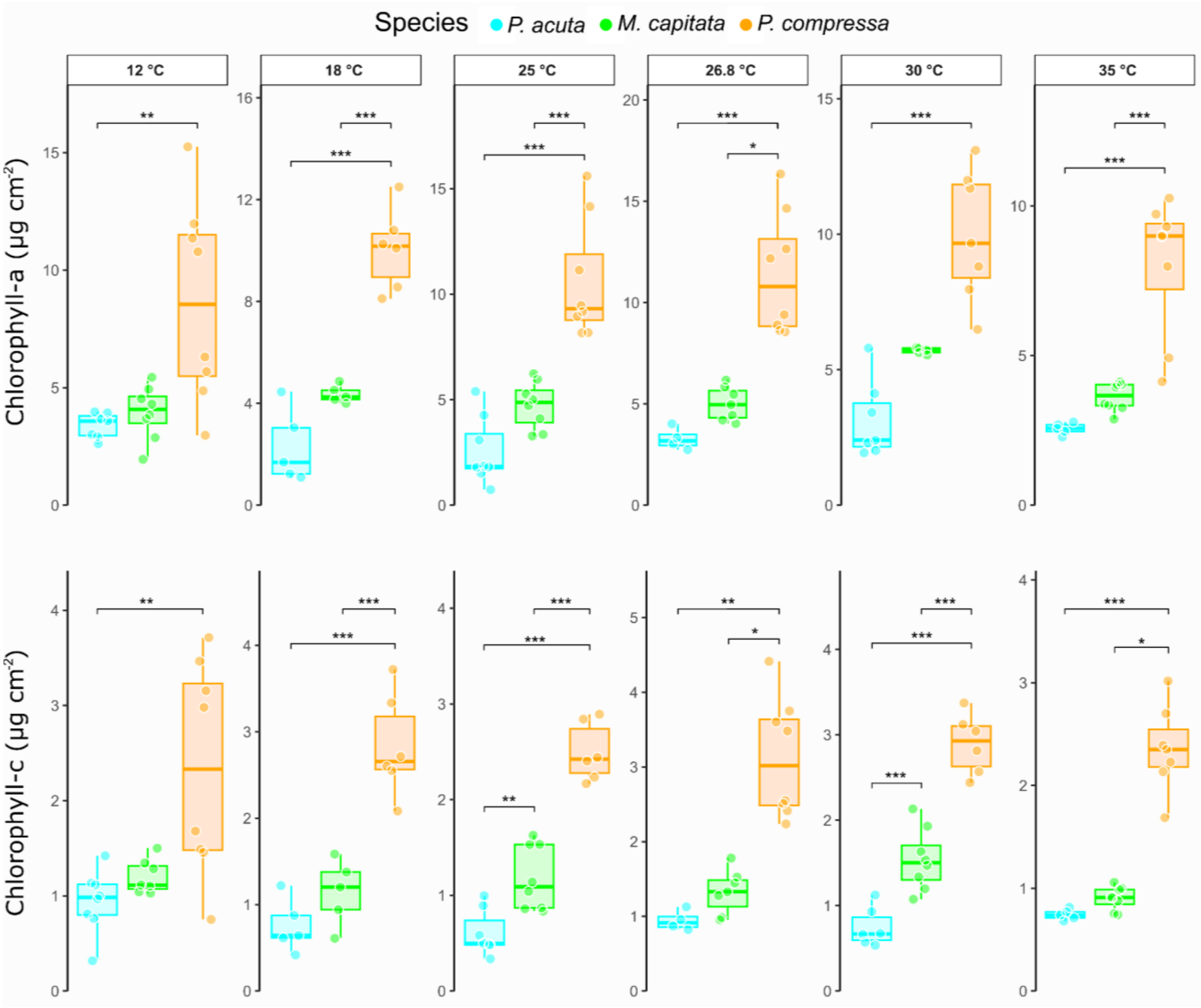


Figure S8. Cross-species chlorophyll concentrations (per host surface area) changes across all temperature treatments. Box plots showing chlorophyll-a (top panels) and chlorophyll-c (bottom panels) concentrations ($\mu g\ cm^{-2}$) per skeleton surface area for *M. capitata* (green), *P. acuta* (cyan), and *P. compressa* (orange) across the six temperature treatments (12°C, 18°C, 25°C, 26.8°C, 30°C, and 35°C). Each box represents the interquartile range with median values indicated by horizontal lines within boxes. Whiskers extend to the most extreme data points within 1.5 times the interquartile range, and individual data points (samples) are overlaid as circles. Horizontal brackets with asterisks indicate statistically significant differences between species pairs (*$p < 0.05$, **$p < 0.01$, ***$p < 0.001$; see Tables S6 and S7 for statistical tests details).

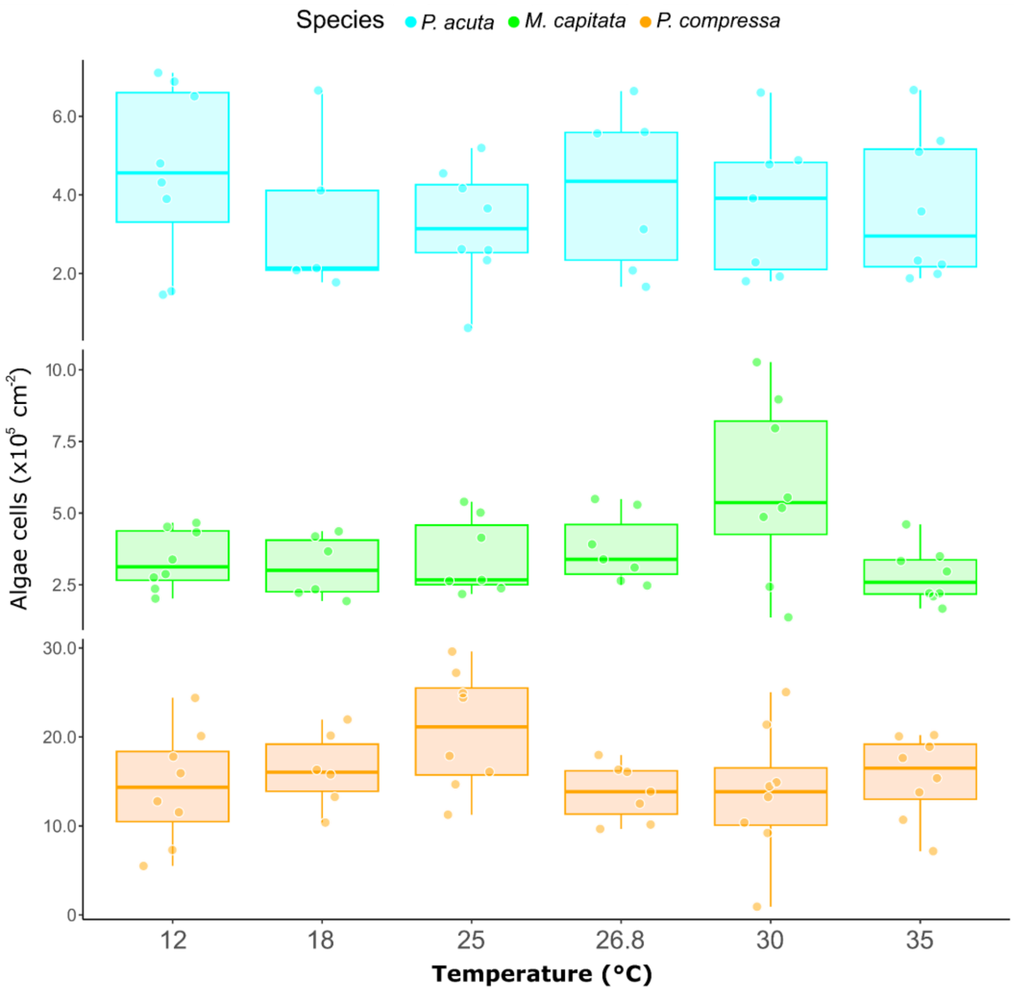


Figure S9. Within-species algae density changes across all temperature treatments. Box plots showing algae cell density per skeleton surface area ($\times10^5$ cells $cm^{-2}$) for *M. capitata* (green), *P. acuta* (cyan), and *P. compressa* (orange) across the six temperature treatments (12°C, 18°C, 25°C, 26.8°C, 30°C, and 35°C). Each box represents the interquartile range with median values indicated by horizontal lines within boxes. Whiskers extend to the most extreme data points within 1.5 times the interquartile range, and individual data points (samples) are overlaid as circles. See Tables S6 and S7 for statistical tests details.

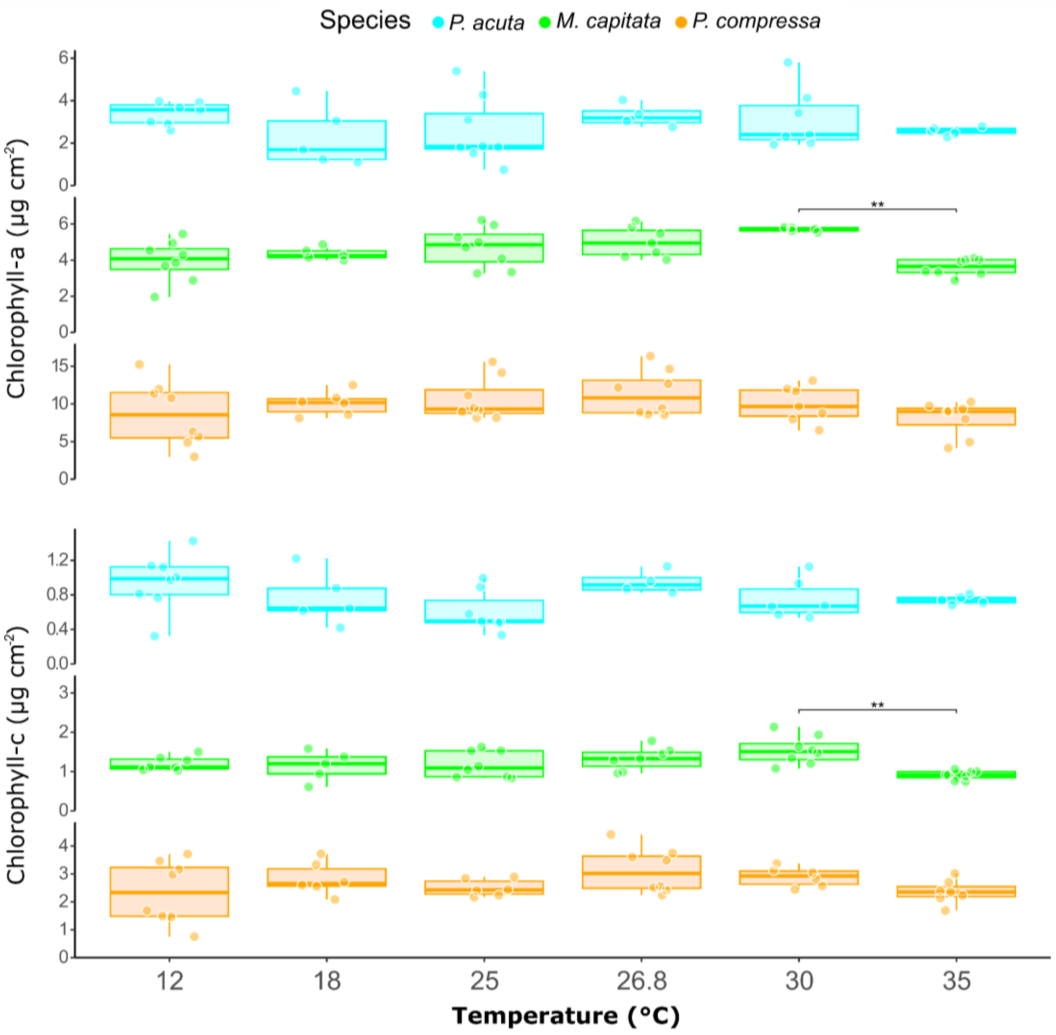


Figure S10. Within-species chlorophyll concentration (per host surface area) changes across all temperature treatments. Box plots showing chlorophyll-a (top panels) and chlorophyll-c (bottom panels) (µg cm$^{-2}$) per skeleton surface area for *M. capitata* (green), *P. acuta* (cyan), and *P. compressa* (orange) across the six temperature treatments (12°C, 18°C, 25°C, 26.8°C, 30°C, and 35°C). Each box represents the interquartile range with median values indicated by horizontal lines within boxes. Whiskers extend to the most extreme data points within 1.5 times the interquartile range, and individual data points (samples) are overlaid as circles. Horizontal brackets with asterisks indicate statistically significant differences between species pairs (**$p < 0.01$; see Tables S6 and S7 for statistical tests details).

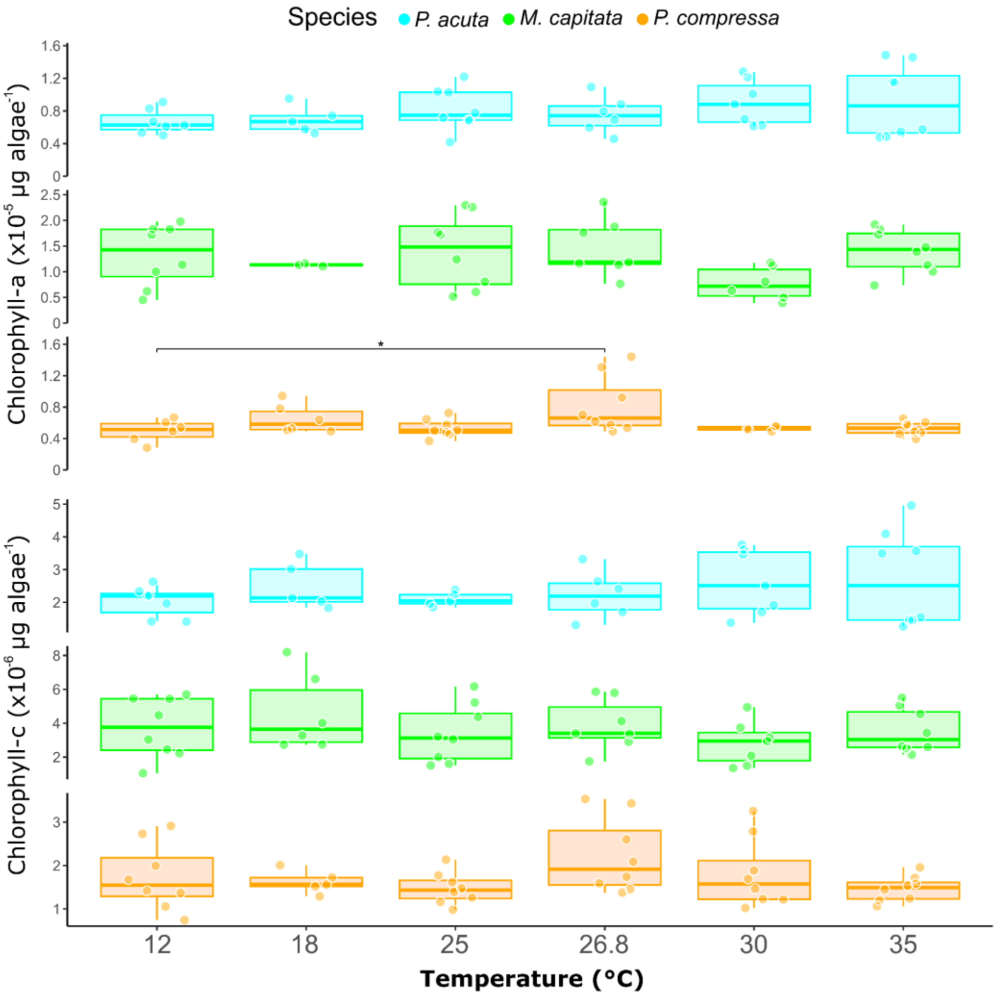


Figure S11. Within-species chlorophyll concentration (per algae cell) changes across all temperature treatments. Box plots showing chlorophyll-a (top panels) and chlorophyll-c (bottom panels) concentrations ($\times 10^{-6}$ µg algae cell$^{-1}$) for *M. capitata* (green), *P. acuta* (cyan), and *P. compressa* (orange) across the six temperature treatments (12°C, 18°C, 25°C, 26.8°C, 30°C, and 35°C). Each box represents the interquartile range with median values indicated by horizontal lines within boxes. Whiskers extend to the most extreme data points within 1.5 times the interquartile range, and individual data points (samples) are overlaid as circles. Horizontal brackets with asterisks indicate statistically significant differences between species pairs (*$p < 0.05$; see Tables S6 and S7 for statistical tests details).

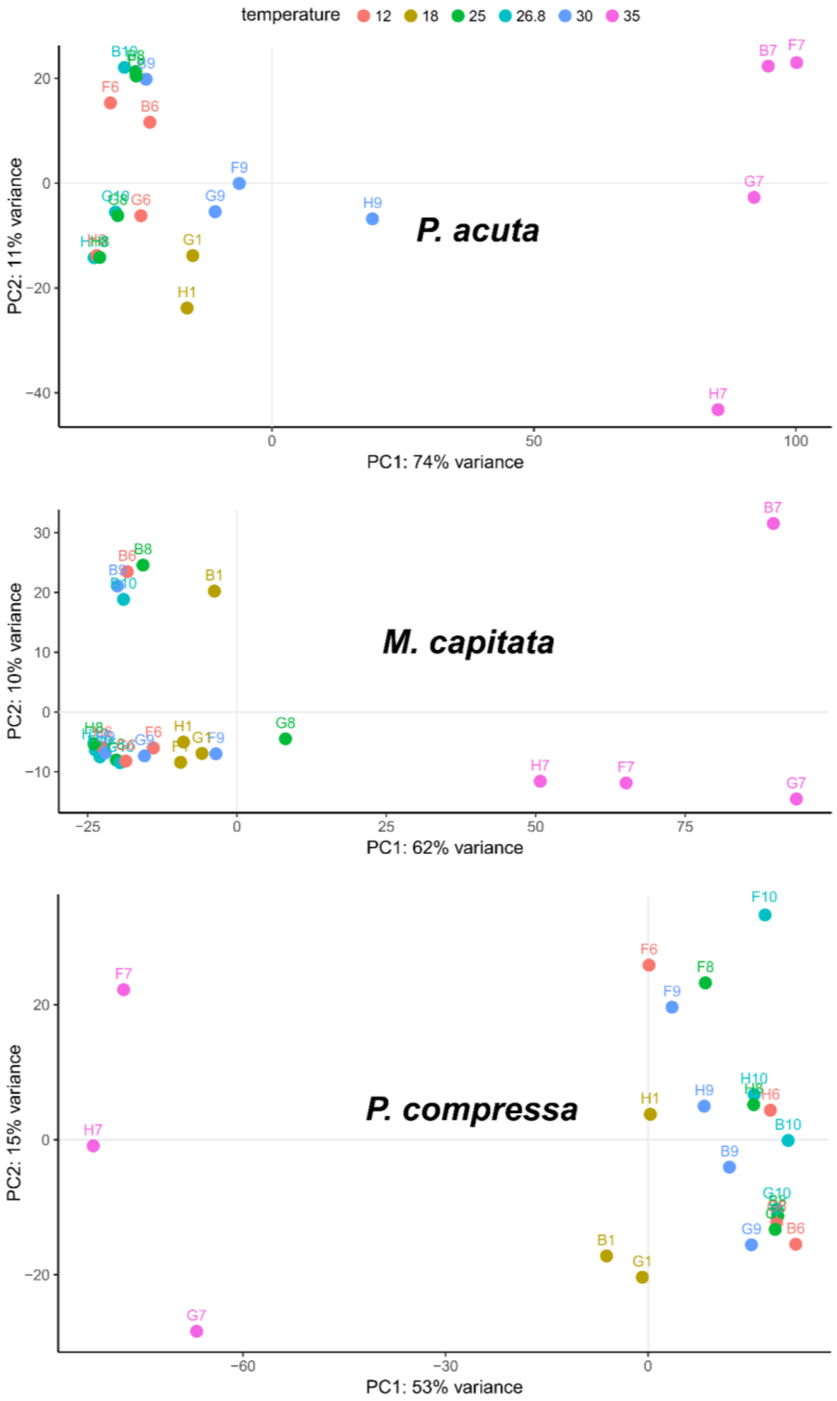


Figure S12. PCA plots of RNA-seq sample-to-sample distances for each coral species. Filtered and normalized gene counts are shown for *M. capitata*, *P. acuta*, and *P. compressa*. Each point represents an individual sample, with colors indicating temperature treatments. Sample labels correspond to coral fragment number and genotype of origin (e.g., B, F, G, H). The first two principal components explain 74%, 62%, and 53% of the total variance for *M. capitata*, *P. acuta*, and *P. compressa*, respectively. The proportion of variance captured is given as a percentage for both the first and second principal components (PC1 and PC2).

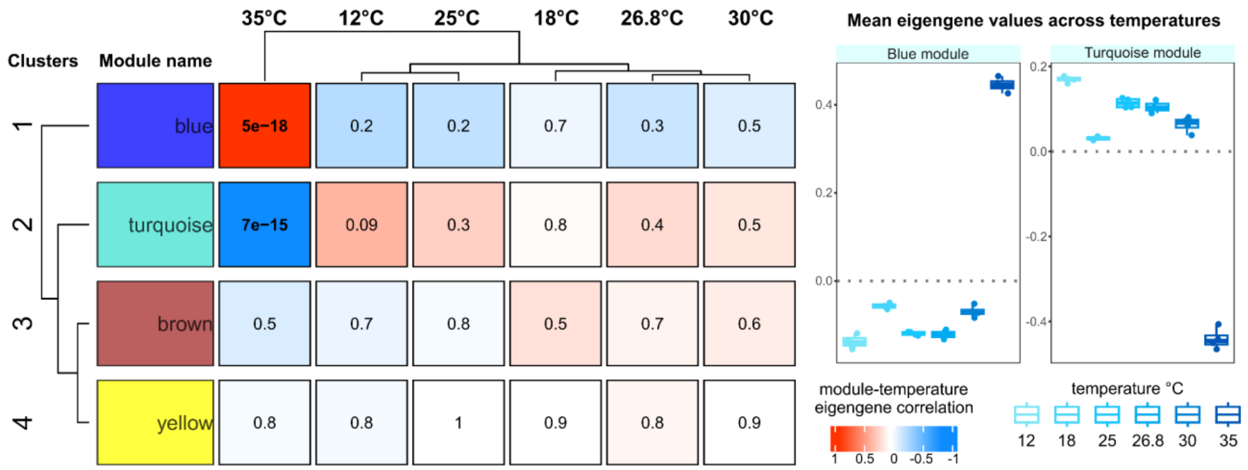


Figure S13. WGCNA results for *P. acuta*. The left panel shows a heatmap of module-temperature correlations with corresponding p-values for each of the identified gene co-expression modules across the six temperature treatments (12°C, 18°C, 25°C, 26.8°C, 30°C, 35°C). Values of correlation between each module and the different clustered temperature treatments range from -1 (anti-correlation) to + 1 (positive correlation). Statistically significant ($p < 0.05$) correlation values are shown in bold. Right panels show boxplots of cross-temperature mean eigengene expression value of modules with significant correlation with at least one temperature treatment. Each dot in the boxplots represents one sample.

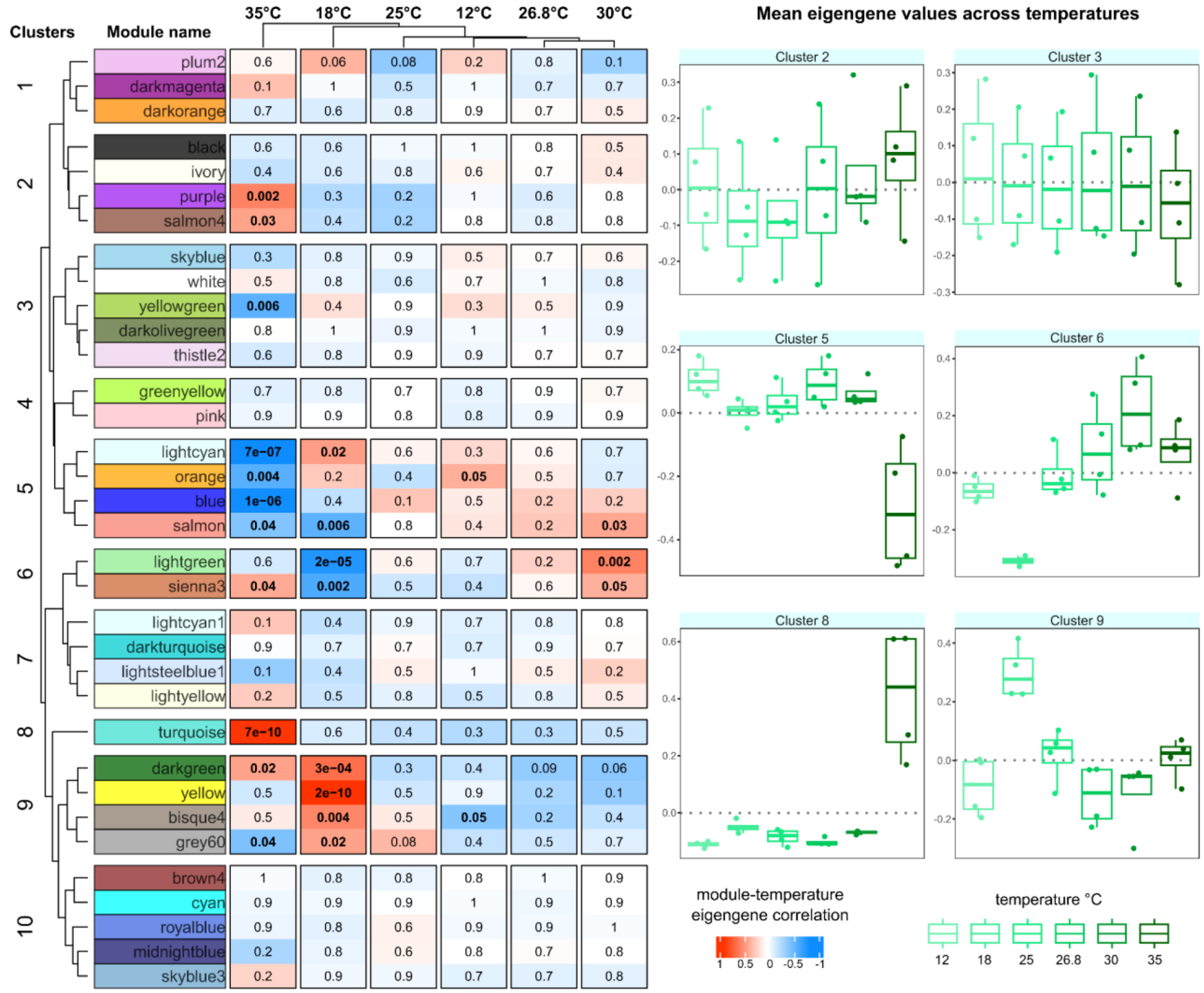


Figure S14. WGCNA results for *M. capitata*. Left panel shows a heatmap of module-temperature correlations with corresponding p-values for each of the identified gene co-expression modules (grouped into 10 clusters) across the six temperature treatments (12°C, 18°C, 25°C, 26.8°C, 30°C, 35°C). Values of correlation between each module and the different clustered temperature treatments range from -1 (anti-correlation) to + 1 (positive correlation). Statistically significant ($p < 0.05$) correlation values are shown in bold. Right panels show boxplots of cross-temperature mean eigengene expression value of clusters with significant correlation with at least one temperature treatment. Each dot in the boxplots represents one sample.

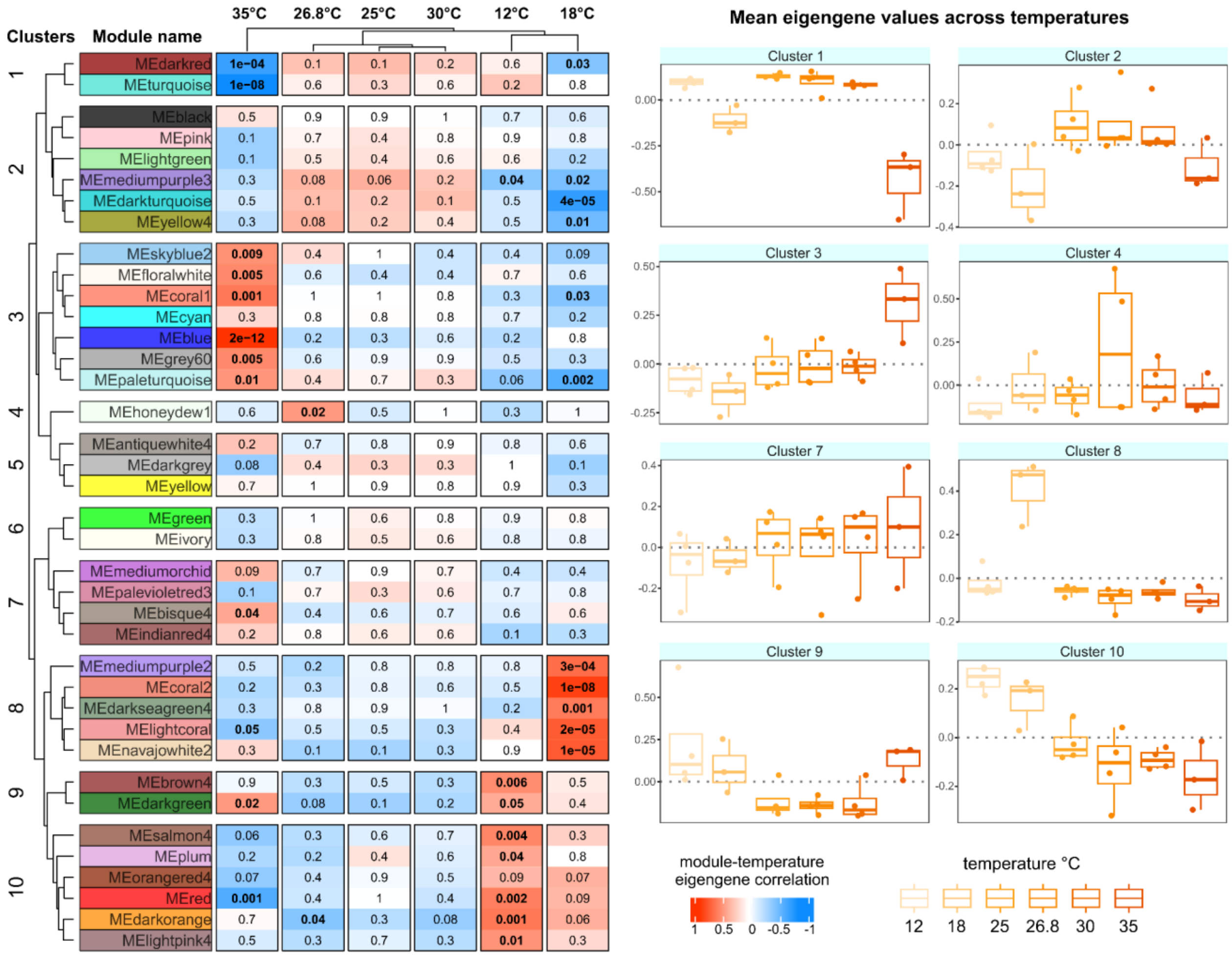


Figure S15. WGCNA results for *P. compressa*. Left panel shows a heatmap of module-temperature correlations with corresponding p-values for each of the identified gene co-expression modules (grouped into 10 clusters) across the six temperature treatments (12°C, 18°C, 25°C, 26.8°C, 30°C, 35°C). Values of correlation between each module and the different clustered temperature treatments range from -1 (anti-correlation) to + 1 (positive correlation). Statistically significant ($p < 0.05$) correlation values are shown in bold. Right panels show boxplots of cross-temperature mean eigengene expression value of clusters with significant correlation with at least one temperature treatment. Each dot in the boxplots represents one sample.

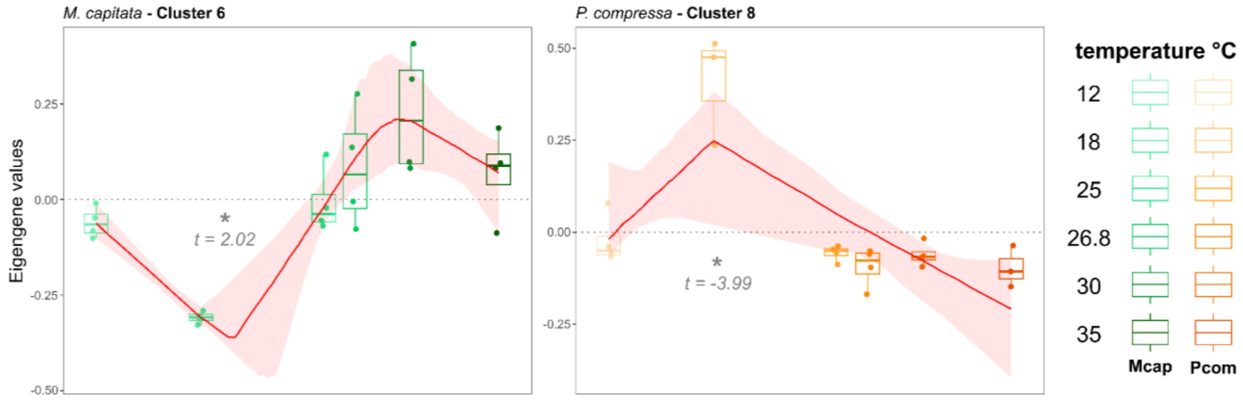


Figure S16. WGCNA clusters with significant break-points at lower temperatures than control. The variation in eigengene expression of clusters with significant temperature correlations was analyzed using regression models to detect break-points (change-points) in the relationship between temperature and the eigengene value. Clusters with significant break-points (t-statistic <-2 or >2) below the control temperature, such as cluster 6 for *M. capitata* and cluster 8 for *P. compressa* were not further analyzed. The red line is the median of the bootstrapped regression fit at each temperature, and shaded red areas indicate the bootstrap confidence intervals around the segmented regression fit.

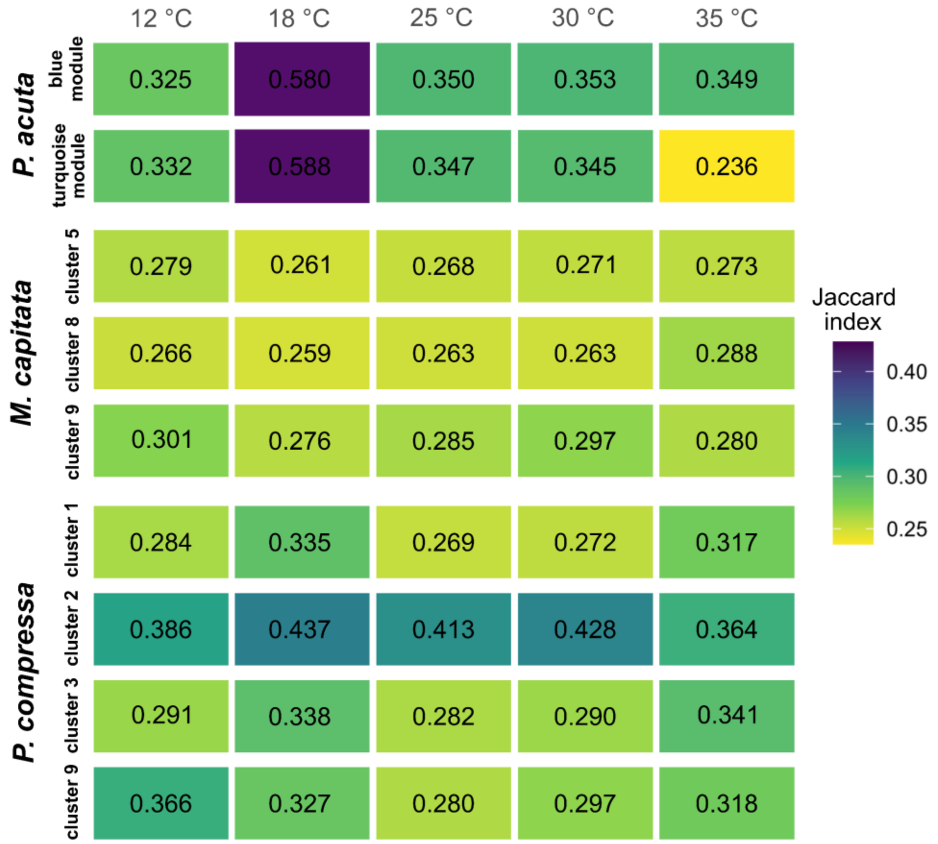


Figure S17. Heatmap showing Jaccard similarity indices for the gene networks in the clusters/modules in Fig. 3A between control temperature (26.8°C) and all other temperature treatments (12°C, 18°C, 25°C, 30°C, 35°C) for each coral species. Each cell represents the Jaccard index (0-1 scale, 1 = identical networks, 0 = no shared network edges) measuring the proportion of shared network edges between cluster/modules at control versus treatment temperatures, with lower values (yellow-green) indicating greater network reorganization and higher values (purple) indicating greater module stability.

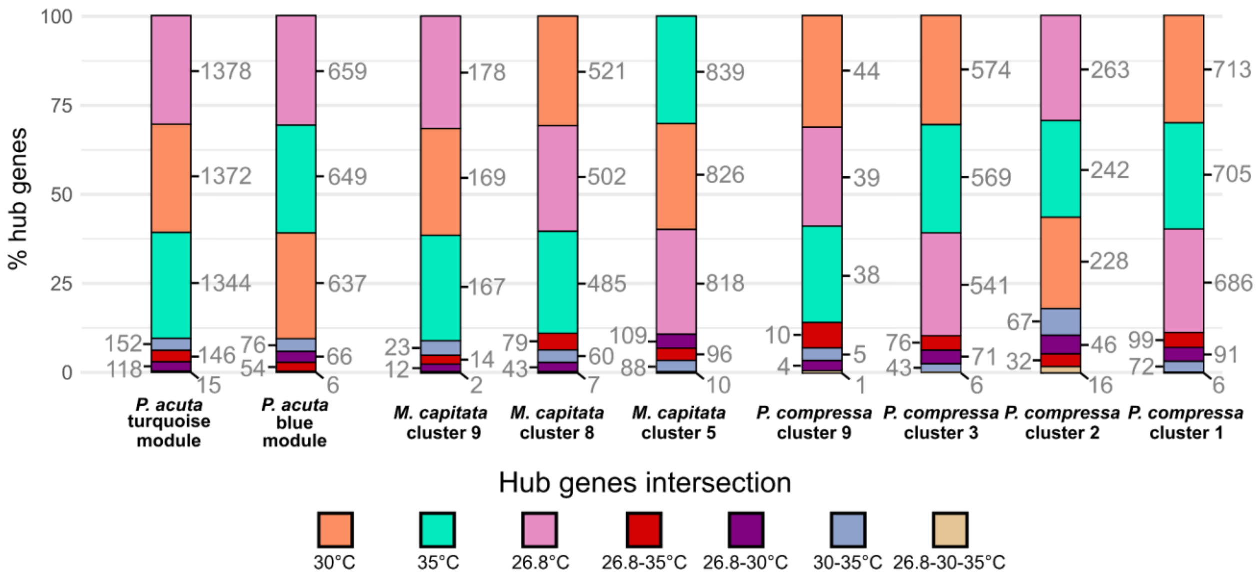


Figure S18. Stacked bar plots showing the intersection patterns of hub genes within significant WGCNA clusters/modules at the control temperature and high (30°C and 35°C) temperature treatments for *P. acuta*, *M. capitata*, and *P. compressa*. Each bar represents 100% of hub genes in a given cluster/module, with colored segments indicating the proportion (intersection) of genes that are: unique to that temperature, shared between two temperatures, or shared across all three temperatures. Numbers adjacent to bar segments indicate the total count of hub genes at each temperature intersection.

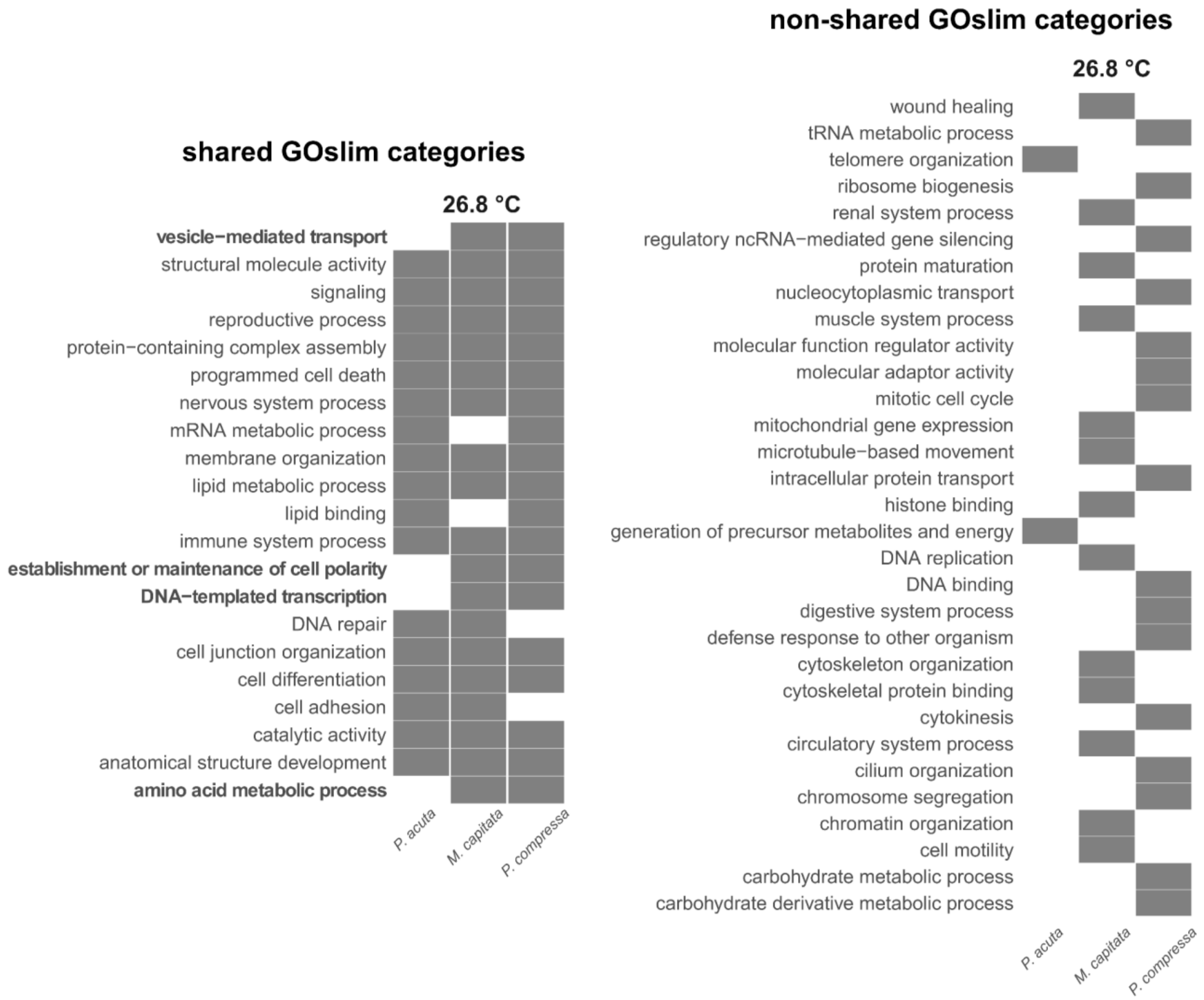


Figure S19. Shared and non-shared enriched GO slim categories across coral species at the control temperature. Enriched GO slim categories among control hub genes from significant WGCNA clusters at the control temperature (26.8°C) for *P. acuta*, *M. capitata* and *P. compressa*. Grey color indicates the baseline level eigengene expression (presence in grey, absence in white).

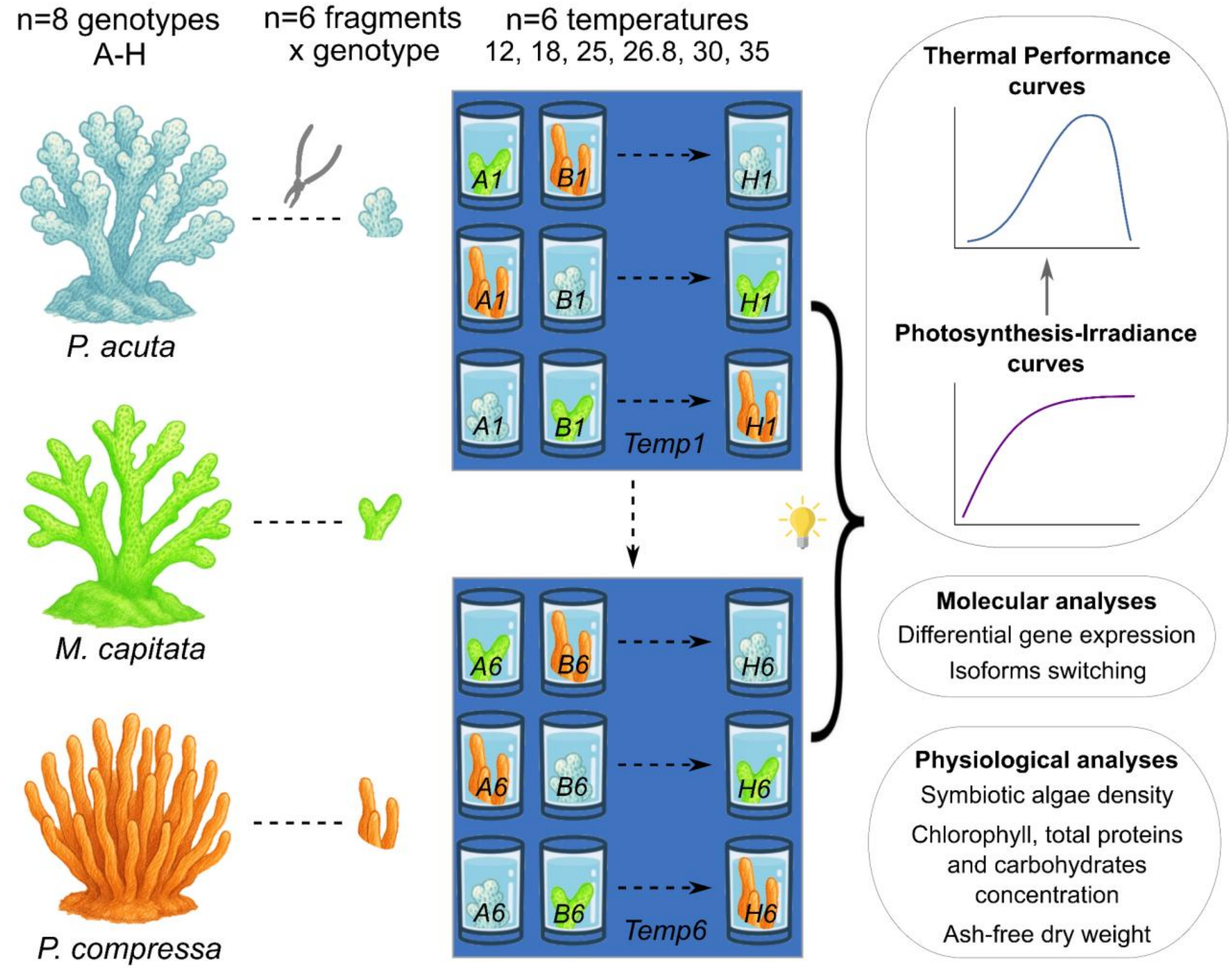


Figure S20. Experimental design overview. For each species, 8 genotypes were collected and cut into fragments (n = 6 fragments per genotype). One fragment per genotype was incubated into one of 6 temperature treatments, and simultaneously exposed to increasing light levels to generate photosynthesis-irradiance curves and thermal performance curves. Following treatment exposure, fragments were collected for physiological and molecular analyses. These include protein, carbohydrate concentration, tissue biomass (for control temperature samples, n=6-8 fragments per species and analysis to assess baseline cross-species differences), chlorophyll concentration and algae density (for all temperatures samples, n=6-8 fragments per temperature per species per analysis, to assess cross-species and within-species differences across temperatures). For molecular analyses, n=3-4 fragments per temperature and species were processed.

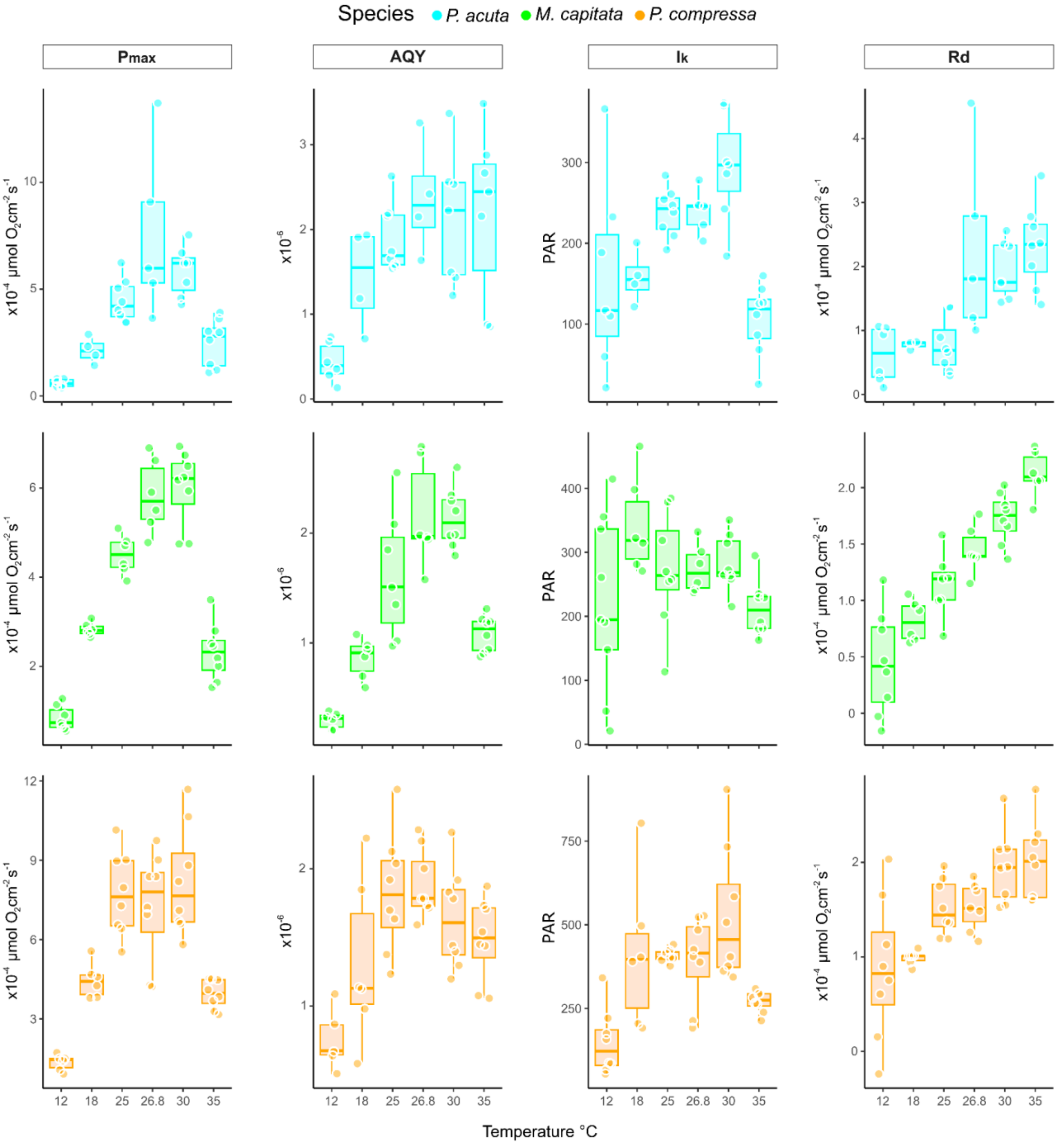


Figure S21. Temperature response of PI-derived parameters across species. Box plots showing the relationship between temperature (°C) and four PI-derived photosynthetic parameters for *M. capitata* (green), *P. acuta* (cyan), and *P. compressa* (orange). Parameters include: Pmax (maximum photosynthetic rate, µmol $O_2$ cm$^{-2}$ s$^{-1}$), AQY (apparent quantum yield), Ik (light saturation parameter, PAR), and Rd (dark respiration rate, µmol $O_2$ cm$^{-2}$ s$^{-1}$). Each box represents the interquartile range with median values indicated by horizontal lines within boxes. Whiskers extend to the most extreme data points within 1.5 times the interquartile range, and individual data points (samples) are overlaid as circles.

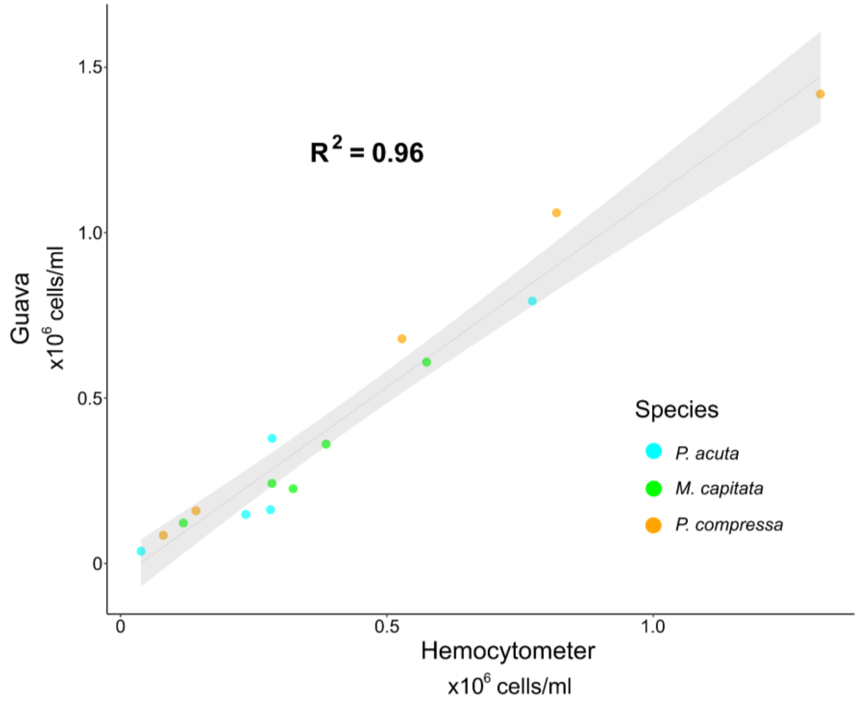


Figure S22. Validation of flow cytometer cell counts against hemocytometer counts. Scatter plot comparing algal cell density measurements obtained using a Guava flow cytometer (×$10^6$ cells/ml) versus traditional hemocytometer counts (×$10^6$ cells/ml) for *M. capitata* (green), *P. acuta* (cyan), and *P. compressa* (orange). Each point represents a paired measurement from the same coral sample. The linear regression line (gray) with 95% confidence interval (shaded area) demonstrates strong concordance between methods ($R^2 = 0.96$).
